# ChIP-seq and RNA-seq for complex and low-abundance tree buds reveal chromatin and expression co-dynamics during sweet cherry bud dormancy

**DOI:** 10.1101/334474

**Authors:** Noémie Vimont, Fu Xiang Quah, David Guillaume-Schöpfer, François Roudier, Elisabeth Dirlewanger, Philip A. Wigge, Bénédicte Wenden, Sandra Cortijo

## Abstract

Chromatin immunoprecipitation-sequencing (ChIP-seq) is a robust technique to study interactions between proteins, such as histones or transcription factors, and DNA. This technique in combination with RNA-sequencing (RNA-seq) is a powerful tool to better understand biological processes in eukaryotes. We developed a combined ChIP-seq and RNA-seq protocol for tree buds (*Prunus avium* L., *Prunus persica* L Batch, *Malus x domestica* Borkh.) that has also been successfully tested on *Arabidopsis thaliana* and *Saccharomyces cerevisiae*. Tree buds contain phenolic compounds that negatively interfere with ChIP and RNA extraction. In addition to solving this problem, our protocol is optimised to work on small amounts of material. Furthermore, one of the advantages of this protocol is that samples for ChIP-seq are cross-linked after flash freezing, making it possible to work on trees growing in the field and to perform ChIP-seq and RNA-seq on the same starting material. Focusing on dormant buds in sweet cherry, we explored the link between expression level and H3K4me3 enrichment for all genes, including a strong correlation between H3K4me3 enrichment at the *DORMANCY-ASSOCIATED MADS-box 5* (*PavDAM5*) loci and its expression pattern. This protocol will allow analysis of chromatin and transcriptomic dynamics in tree buds, notably during its development and response to the environment.

## BACKGROUND

The term ‘epigenetics’ has traditionally been used to refer to heritable changes in gene expression that take place without altering DNA sequence (Wolffe and Matzke, 1999), but it is also used, in a broader sense, to refer to modifications of the chromatin environment (Miozzo et al., 2015). Epigenetic modifications are important for a wide range of processes in plants, including seed germination (Nakabayashi et al., 2005), root growth (Krichevsky et al., 2009), flowering time (He et al., 2003), disease resistance (Stokes et al., 2002) and abiotic stress responses (Zhu et al., 2008). Post-transcriptional modifications of histone proteins and chromatin structure regulate the ability of transcription factors (TFs) to bind DNA and thereby influence gene expression (Lee et al., 1993; Narlikar et al., 2002). Analysing the dynamics of chromatin modifications and DNA-protein interactions is a critical step to fully understand how gene expression is regulated. Chromatin Immunoprecipitation (ChIP) is one of the few methods enabling the exploration of *in vivo* interactions between DNA and proteins such as histones and TFs. When followed by next generation sequencing (ChIP-seq), this method allows the detection of these interactions at a genome-wide scale. Since chromatin modifications and the regulation of gene expression are tightly linked, ChIP-seq for chromatin marks and TFs are often combined with RNA-sequencing (RNA-seq) to extract key features of the role of chromatin modification and TF binding in regulating transcription. While ChIP-seq is routinely performed in plant model organisms like *Arabidopsis thaliana*, it is still a challenge to carry it out on tree buds. The numerous ChIP protocols published in plants and mammals [(Cortijo et al., 2018; Kaufmann et al., 2010; Li et al., 2014; Nelson et al., 2006; Ricardi et al., 2010; Saleh et al., 2008; Wal and Pugh, 2012; Xie and Presting, 2016; Yamaguchi et al., 2014) to cite just some] cannot be directly used for plant materials with high phenolic content (alkaloids or lignified cell walls) like tree buds (Bílková et al., 1999). Such chemicals need to be chelated during the chromatin extraction to prevent inhibition of downstream processes. In addition, the requirement for large amounts of starting material in previous protocols has made it a challenge to perform ChIP on low-abundance tissues such as tree buds. Here we present an efficient protocol for ChIP-seq and RNA-seq on complex, low-abundance tree buds, with the possibility of studying trees growing in the field and rapid chromatin dynamics owing to an improved cross-linking method (Figure 1). While several studies include ChIP-seq performed in trees, some with improvements described in this protocol such as using a chelator of interfering compounds or performing cross-linking in frozen material (Hussey et al., 2015; de la Fuente et al., 2015; Leida et al., 2012), this is the first step-by-step detailed ChIP-seq protocol in trees that includes all of these improvements:

- Our ChIP-seq/RNA-seq protocol can be carried out on complex plant tissues that contain interfering compounds (phenolic complexes, scales, protective layers), by adding chelators of these compounds in the extraction buffers.
- The cross-linking step is performed on frozen, pulverised material, thus allowing sample collection in the field, where cross-linking equipment is not available. It also allows studying fast responses by flash freezing material immediately after a stimulus (e.g. transient temperature stress) rather than cross-linking directly on fresh tissue or cells. In previous protocols, the cross-linking step was performed using a vacuum and lasted at least 10 minutes and up to 1 hour (Kaufmann et al., 2010; Ricardi et al., 2010; Saleh et al., 2008; Xie and Presting, 2016). Moreover, cross-linking on powder allows for a more homogenous cross-linking, as it is almost impossible to have a homogenous penetration of the formaldehyde in tree buds that are protected by an impermeable and rigid lignin rich wall (Bílková et al., 1999).
- By using frozen, pulverized material, ChIP-seq and RNA-seq can be performed on the same starting material for a direct and robust comparison of epigenetic regulation and gene expression.
- Our protocol can be used to perform ChIP-seq and RNA-seq on a small amount of biological material. We optimised this protocol to start from 0.2 to 0.5 g of buds, which is considerably lower than the usual amount of 0.8 to 5 g of starting material for ChIP protocols in plant tissues (Haring et al., 2007; Kaufmann et al., 2010; Leida et al., 2012; Ricardi et al., 2010; Saleh et al., 2008; Xie and Presting, 2016), or the 4g of tree buds previously used (de la Fuente et al., 2015; Leida et al., 2012).

**Figure 1.**
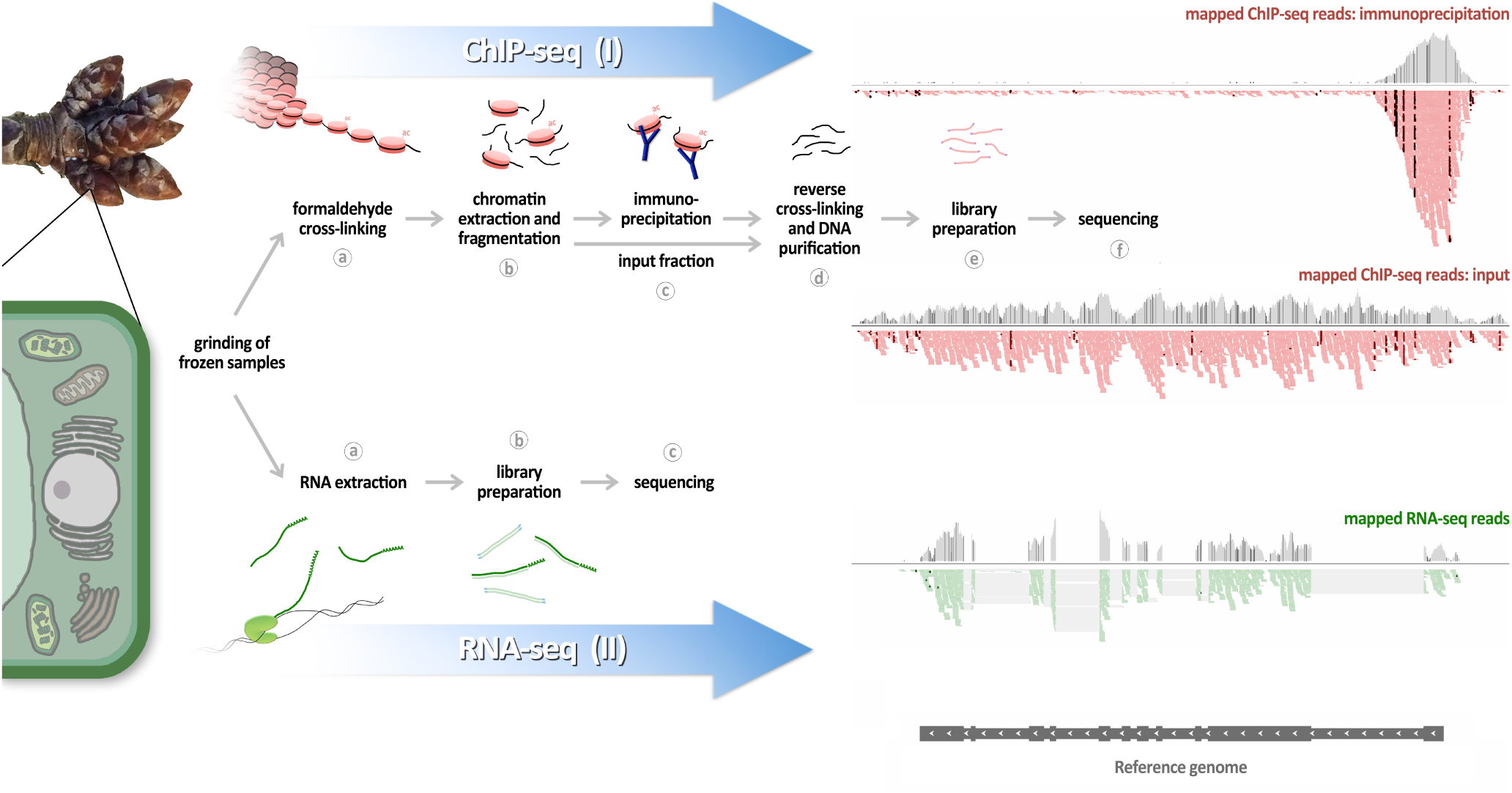
Workflow. Outline of the two main modules of the protocol: (I) ChIP-seq (top) and (II) RNA-seq (bottom). Each module starts with the same biological material (ground frozen material). (I) (a) The ChIP-seq module starts with a cross-linking step on frozen powder to stabilise interactions between DNA and proteins. (b) The chromatin is extracted using different buffers and then fragmented by sonication or MNase digestion. (c) Proteins of interest, among the protein/DNA complexes, are immunoprecipitated using specific antibodies coupled to magnetic beads. An aliquot of chromatin is set aside as an input fraction. (d) After different wash steps, a reverse cross-linking step is performed, and the DNA is isolated using SPRI beads. (e) The purified DNA is used in library preparation, and (f) is then sequenced. (II) (a) The RNA-seq module starts with RNA extraction from the frozen powder. (b) This DNAse treated RNA is then used in library preparation, and (c) sequenced.

We have used this protocol to analyse histone modification profiles in several tree species. In a first instance, we analysed H3K27me3 in buds of *Prunus persica* L Batch (peach) and could replicate previously published results (de la Fuente et al., 2015). To demonstrate the versatility of this protocol, we also successfully performed ChIP-seq for H3K27me3 and H3K4me3 in sweet cherry *(Prunus avium* L.) and apple (*Malus x domestica* Borkh.). Furthermore, we directly compared expression level and enrichment for H3K4me3 by performing ChIP-seq for H3K4me3 and RNA-seq on the same sweet cherry bud samples. We demonstrated the correlation between chromatin status and gene expression for *AGAMOUS* (*AG*) and *ELONGATION FACTOR 1* (*EF1*) that are known to be under control of H3K27me3 and H3K4me3, respectively (Saito et al., 2015). We expended our analysis of chromatin and expression in cherry buds harvested at different stages of dormancy, first to *DORMANCY-ASSOCIATED MADS-box 6 and 5* (*PavDAM6* and *PavDAM5*) genes, which are key regulators of dormancy in trees, and then to the entire genome. Dormancy is an important developmental stage of fruit trees and is characterised by a period of repressed growth that allows trees to persist under low winter temperature and short photoperiod (Faust et al., 1997). A proper regulation of the timing of the onset and release of bud dormancy is crucial to ensure optimal flowering and fruit production in trees. Consequently, unravelling the associated molecular mechanisms is essential and numerous studies have been conducted in trees to answer this question. De la Fuente and colleagues (de la Fuente et al., 2015) have shown that *DORMANCY-ASSOCIATED MADS-box* (*DAM*)-related genes are up-regulated in dormant peach buds. These genes are involved in the regulation of bud dormancy under unfavorable climatic conditions in peach (de la Fuente et al., 2015; Leida et al., 2012; Yamane et al., 2011), leafy spurge (Horvath et al., 2010), pear (Saito et al., 2015), apple (Mimida et al., 2015) and apricot (Sasaki et al., 2011). We find that dormancy-associated *PavDAM6* and *PavDAM5* genes are more expressed in dormant buds than in non-dormant buds and that H3K4me3 occupancy is associated with *PavDAM5* expression level. We also find significant changes in H3K4me3 level during dormancy for 671 genes, and that these changes are positively associated with transcriptional changes during dormancy. Our results show the potential for future exploration of the link between chromatin dynamics and expression at a genome-wide level during tree bud dormancy. Moreover, our combined ChIP-seq and RNA-seq protocol, which is working on many tree species, will allow a better understanding of transcriptional regulatory events and epigenomic mechanisms in tree buds.

## RESULTS

### Validation of the protocol robustness in peach

In order to validate our protocol, we analysed H3K27me3 enrichment in the gene body of a *DAM* gene cluster on peach (*Prunus Persica* L Batch) non-dormant buds (Figure 2). H3K27me3 enrichment is known to be associated with a repressed transcriptional state. It has been shown that *DORMANCY-ASSOCIATED MADS-box* (*DAM*)-related genes display contrasting H3K27me3 profiles in dormant and non-dormant buds, with H3K27me3 signal only observed in non-dormant buds (de la Fuente et al., 2015). We observe a higher enrichment for H3K27me3 at the peach *PpeDAM5* and *PpeDAM6* loci compared with the *PpeDAM3* gene in non-dormant buds, or compared to the gene *EF1,* known to be enriched in H3K4me3 and depleted in H3K27me3 (Figure 2, Figure 3a). The replication of previously published results at the *DAM* genes confirms that our improved ChIP-seq protocol is working on tree buds.

**Figure 2.**
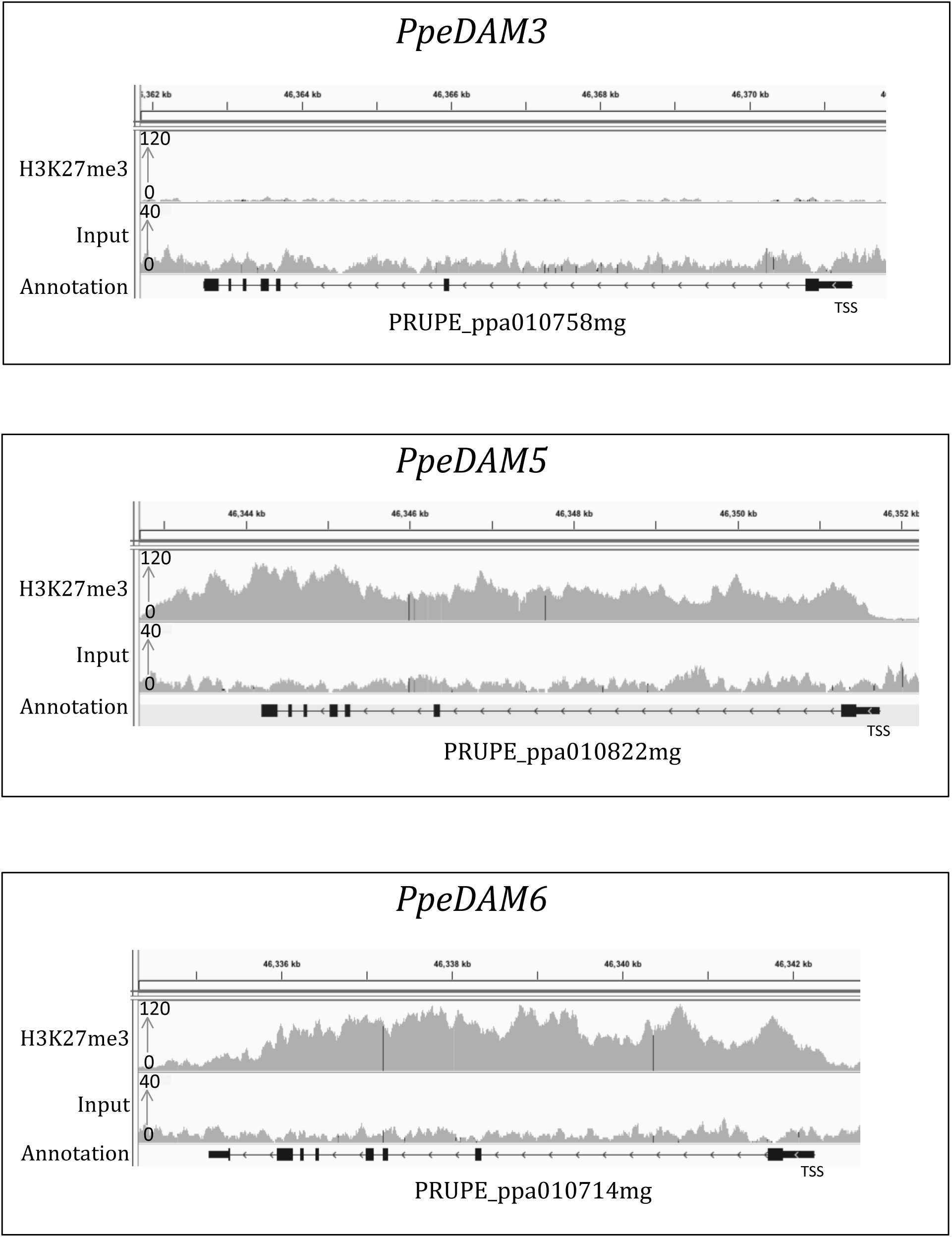
H3K27me3 ChIP-seq profile at the *DAM* gene cluster in peach. IGV screenshot of ChIP-seq data for H3K27me3 and its corresponding input performed on non-dormant peach buds (*Prunus persica*) at PpeDAM3, PpeDAM5 and PpeDAM6 genes. Genes are represented by black lines, with arrows indicating gene directionality and rectangles representing exons.

**Figure 3.**
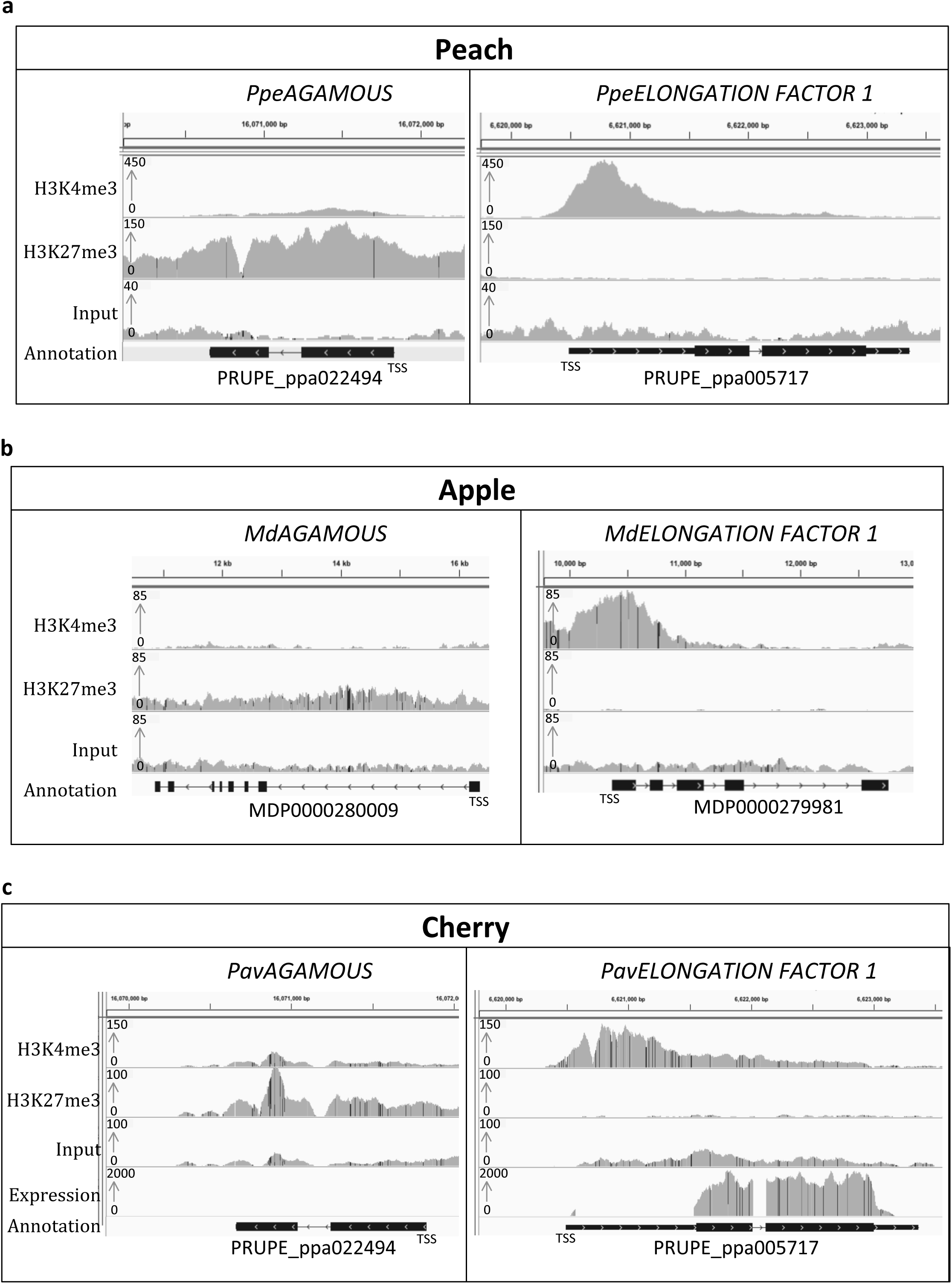
H3K27me3 and H3K4me3 ChIP-seq and RNA-seq profiles three fruit tree species. IGV screenshot of ChIP-seq data for H3K27me3 and H3K4me3 and their corresponding inputs performed on peach (a), apple (b) and sweet cherry buds (c) for two control genes: ELONGATION FACTOR 1 as a positive control of H3K4me3 and AGAMOUS as a positive control of H3K27me3. RNA-seq was also carried out on sweet cherry buds (c). Genes are represented by black lines, with arrows indicating gene directionality and rectangles representing exons.

### Successful application of this protocol on different species

To demonstrate the versatility of this protocol, we performed ChIP-seq for two histone marks (H3K27me3 and H3K4me3) on buds of three tree species: peach, apple (*Malus x domestica* Borkh.) and sweet cherry *(Prunus avium* L.) (Figure 3). We analysed the signal at the genes *ELONGATION FACTOR 1* (*EF1*), known to be enriched in H3K4me3, and *AGAMOUS* (*AG*), known to be enriched in H3K27me3 (Saito et al., 2015). While H3K27me3 is associated with a repressive transcriptional state, H3K4me3 is on the contrary associated with transcriptional activation. We observed a strong H3K4me3 signal at *EF1* and enrichment for H3K27me3 at *AG* locus for the three species (Figure 3). This result confirms that this ChIP-seq protocol works on buds for several tree species. To test the adaptability of this protocol for other biological system, we also performed ChIP-qPCR in *Arabidopsis thaliana* and in *Saccharomyces cerevisiae*. We reproduced previously published results for the histone variant H2A.Z at *HSP70* locus in *Arabidopsis thaliana* (Cortijo et al., 2017; Kumar and Wigge, 2010), and for the binding of the TF Hsf1 at the SSA4 promoter in yeast (Erkina and Erkine, 2006), Suppl. Figure 1). This protocol is thus versatile and can be used on buds for several tree species as well as in other biological systems such as *Arabidopsis thaliana* and yeast.

### Association between gene expression and H3 K27 m 3 enrichment at *PavDAM* loci

To test if chromatin state and gene expression can be directly compared using this protocol, we carried out RNA-seq and ChIP-seq on the same starting material of sweet cherry floral buds. To start with, we analysed expression together with enrichment for H3K27me and H3K4me3 for *PavEF1* and *PavAG* genes. We observe that *PavEF1*, marked by H3K4me3, is highly expressed, while *PavAG*, marked by H3K27me3, is very lowly expressed (Figure 3c, Suppl. Figure 4).

We then compared the abundance of H3K4me3 histone mark to the expression patterns of *PavDAM5* and *PavDAM6* during the dormancy period at three different dates for sweet cherry floral buds (October, December and January; Figure 4). Firstly, we defined if the flower buds harvested are in endodormancy or ecodormancy at each time-point (Figure 4a). The time of dormancy release, after which the buds are in ecodormancy, is defined when the percentage of bud break reaches 50% at BBCH stage 53 (Meier, 2001). We observe that samples harvested in October and December are in endodormancy and the ones harvested in January are in ecodormancy (Figure 4a). *PavDAM5* and *PavDAM6* are two key genes involved in sweet cherry dormancy corresponding to the peach *PpeDAM5* and *PpeDAM6*, respectively. We find that *PavDAM6* is highly expressed in October at the beginning of endodormancy and that its expression decreases in December and January (Figure 4b), and that *PavDAM5* is highly expressed in deep dormancy (December) and less expressed in ecodormancy (January, Figure 4b). These results are in agreement with previous observations showing that *DAM* genes are up-regulated in dormant peach buds (de la Fuente et al., 2015). We observe H3K4me3 enrichment at the beginning of these genes, at the level of the first exon (Figure 4c, Suppl. Figure 5) as expected for this chromatin mark. A low or no enrichment was found for the two controls (H3 and INPUT, Figure 4c) meaning that the enrichment seen for H3K4me3 at these genes is relevant. We observe that H3K4me3 enrichment at *PavDAM5* is higher in December compared to October and January (Figure 4c). This is in agreement with the higher *PavDAM5* expression in December. We observe that H3K4me3 enrichment at *PavDAM6* is also higher in December compared to October and January, while its peak of expression is in October (Figure 4c).

**Figure 4.**
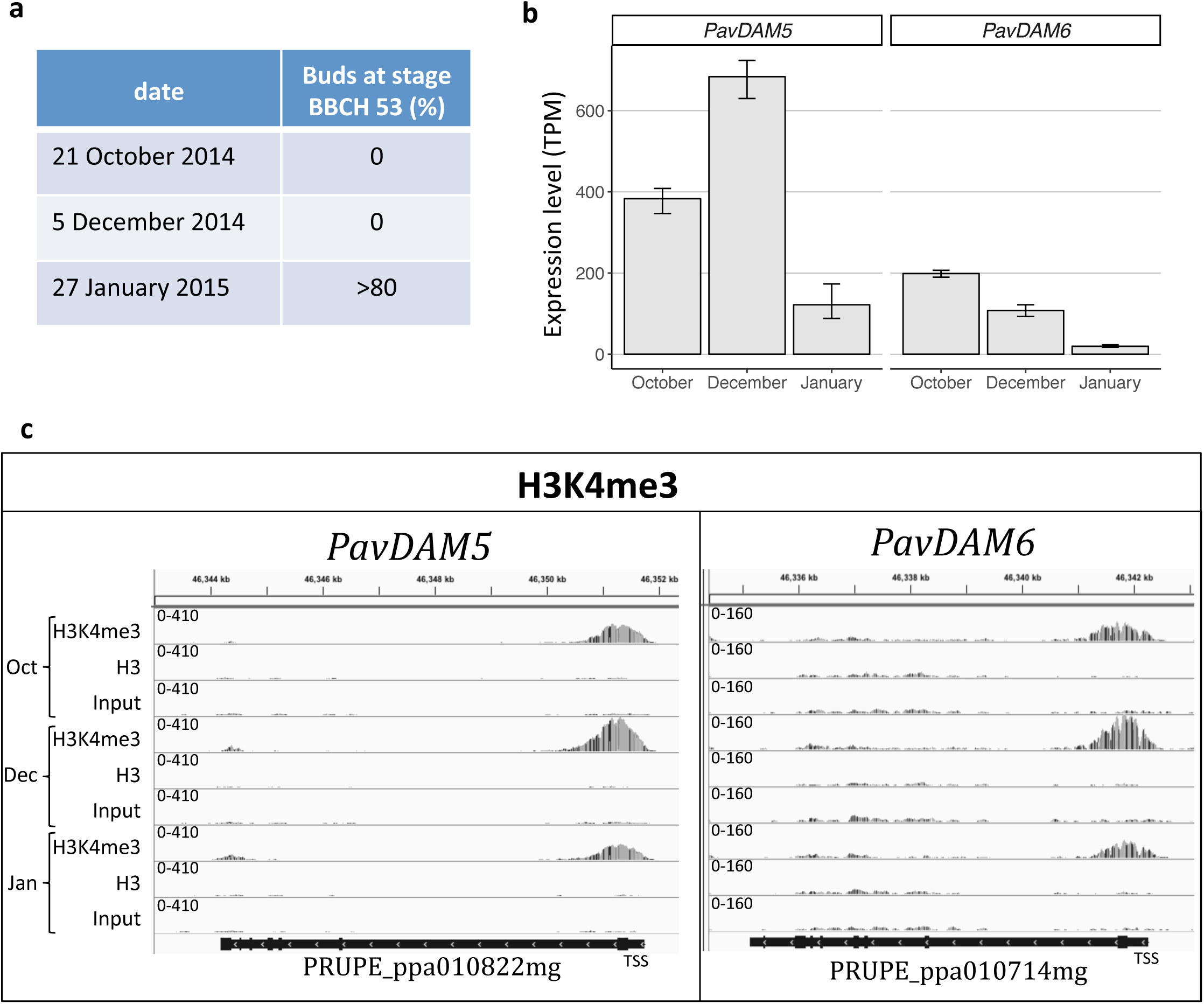
Expression and H3K4me3 profiles of *PavDAM5* and *PavDAM6* genes during dormancy in sweet cherry flower buds. a) Evaluation of bud break percentage under forcing conditions (n=5). b) Transcriptional dynamics of *PavDAM5* and *PavDAM6* genes at three different dates (October 21st 2014, December 5th 2014 and January 27th 2015; n=3). Expressions are represented in TPM (Transcripts Per kilobase Million). C) IGV screenshot of the first biological replicate of ChIP-seq data for H3K4me3 and H3 at three different dates (October 21st 2014, December 5th 2014 and January 27th 2015) and their corresponding inputs at *PavDAM5* and *PavDAM6* loci. Genes are represented by black rectangles, with arrows indicating gene directionality and bigger rectangles representing exons.

### Direct comparison of ChIP-seq and RNA-seq data

To further investigate the link between gene expression and H3K4me3 enrichments during bud dormancy, we analysed genome-wide changes between time-points in expression level as well as H3K4me3 signal. For this, we employed a gene centric approach by measuring the strength of the H3K4me3 signal at each gene, and identifying genes showing significant changes between at least two of the three time-points (see material and methods for more detail). We identified 671 genes that show significant changes in H3K4me3 between at least two of the sampling dates. We then performed a hierarchical clustering of these genes based on a Z-score [(signal for a time-point - average over all time-points)/ STD across all time-points], which normalises for differences in H3K4me3 signal between genes, thus allowing direct gene-by-gene comparisons of their patterns across the studied time-points. We observe a reduction in H3K4me3 signal over time, and in particular between endodormancy and ecodormancy for most genes (Figure 5a, see Suppl. Figure 7bc for examples of H3K4me3 signal at a few genes). We then compared changes in H3K4me3 and expression over time, for the genes in the main clusters depicted in Figure 5b and Figure 5c. Genes in the purple (233 genes) and blue (313 genes) clusters are associated with simultaneous reductions in H3K4me3 signal and expression over time (Figure 5c). Genes in the green (53 genes), the gold (40 genes) and the orange (13 genes) clusters show increases in H3K4me3 signal and in expression over time (Figure 5c). Genes in the red cluster (19 genes) are characterised by a transient reduction in H3K4me3 signal and expression in December (Figure 5c).

**Figure 5.**
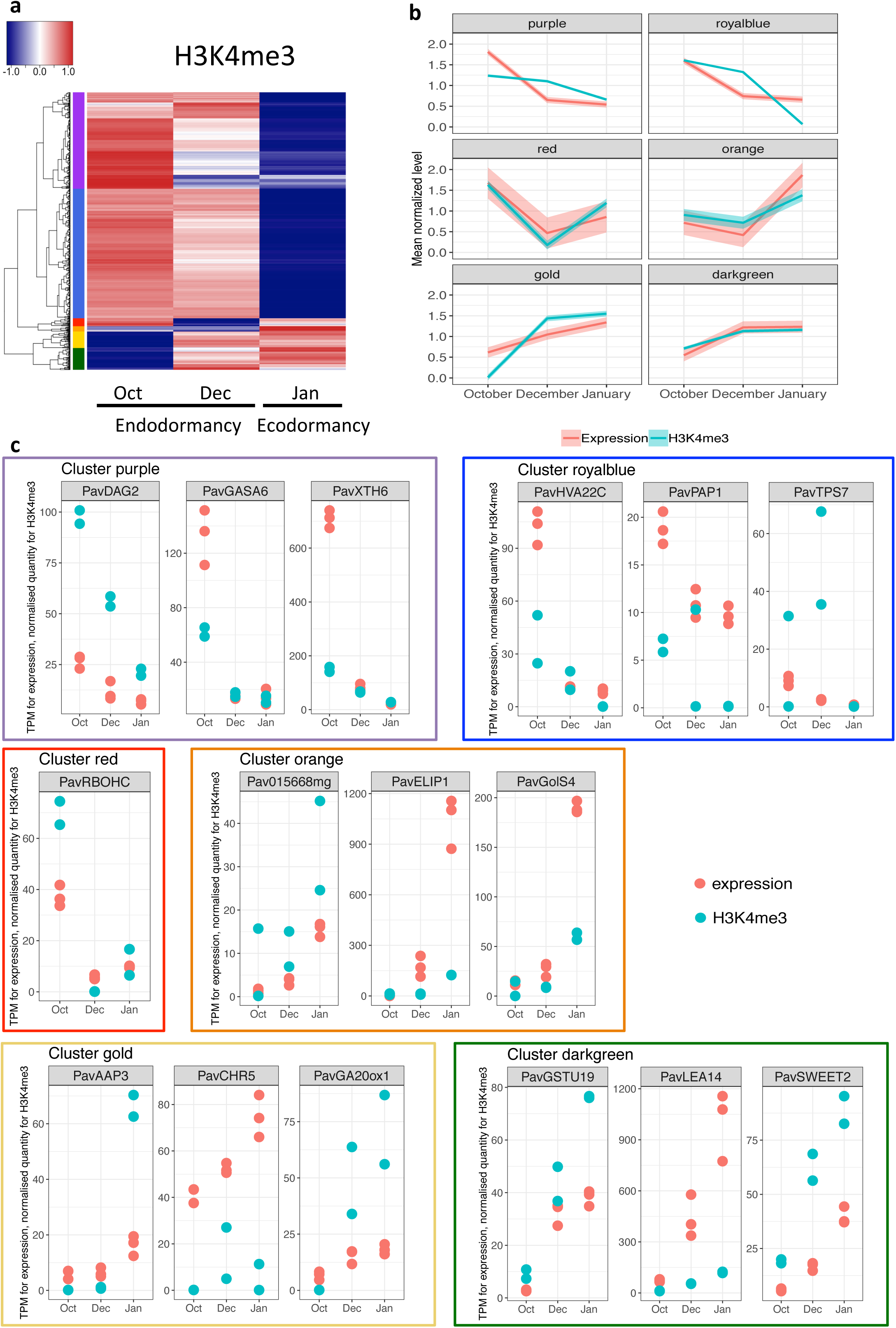
Genome wide comparison of H3K4me3 and expression in sweet cherry flower buds in endodormancy and ecodormancy. a) Hierarchical clustering of genes showing a change in H3K4me3 level between the time points, based on the Z-score of the normalised H3K4me3 level around the TSS of each gene [Z-score = (signal for a time point - average over all time points)/ STD across all time points]. The result is represented as a heatmap where red indicates a high Z-score and blue a low Z-score. The genes were separated into six clusters, indicated by the side coloured bar. b) Average signal in each time point for genes in clusters showing changes in H3K4me3 over time (identified in figure 5a) for expression (red) and H3K4me3 (blue). Mean normalised expression and H3K4me3 levels was used (level in one time-point/ average across the entire time course for a given gene). The standard deviation is also shown as a transparent ribbon. c) Expression level (TPM) and normalised H3K4me3 signal over time for a few individual genes in each of the 6 clusters identified on Figure 5a. Each point represents one biological replicate. Normalised H3K4me3 signal corresponds to the number of H3K4me3 ChIP reads in a 2000bp window around the TSS of each gene normalised by the number of H3 ChIP reads in the same window.

We explore signalling pathways that were represented in the different clusters (Figure 5c, Suppl. figure 7b, Suppl. Table 2). Among the genes classed in the green, gold and orange clusters, that increase expression over time, we identified the *GLUTATHION S-TRANSFERASE19* (*PavGSTU19*), *LATE EMBRYOGENESIS ABUNDANT14* (*PavLEA14*) and GALACTINOL SYNTHASE 4 (*PavGolS4*) genes (Figure 5c), potentially involved in the response to drought and oxidative stresses (Hara et a., 2010; Nishizawa et al., 2008; Singh et al., 2005), together with genes associated with growth and cellular activity such as *PavSWEET2*, *GIBBERELLIN 20 OXIDASE 1* (*PavGA20ox1*) and *AMINO ACID PERMEASE3* (*PavAAP3*). On the other hand, genes from clusters purple and blue that are downregulated during ecodormancy include *XYLOGLUCAN ENDOTRANSGLUCOSYLASE/HYDROLASE6* (*PavXTH6*), *DOF AFFECTING GERMINATION2* (*PavDAG2*), *GA-STIMULATED IN ARABIDOPSIS6* (*PavGASA6*), *TREHALOSE-PHOSPHATASE/SYNTHASE7* (*PavTPS7*), *PavHVA22C*, and *PHYTOCHROME-ASSOCIATED PROTEIN1* (*PavPAP1*) (Figure 5c). Homologs of *XTH6*, *DAG2*, *GASA6* and *HVA22C* were found to be activated under cold, drought and other stress-associated stimulus while *TPS7* and *PAP1* are more likely associated with metabolism and auxin pathway. We also found *RESPIRATORY BURST OXIDASE HOMOLOG C* (*PavRBOHC*) in the cluster red, transiently downregulated in December, of which the homolog in Arabidopsis is involved in reactive oxygen species production.

## DISCUSSION

### ChIP-seq and RNA-seq protocol: new epigenetic perspectives in trees

In this study, we described a combined ChIP/RNA-seq protocol for low abundance and complex plant tissues such as tree buds. This method allows a robust comparison of epigenetic regulation and gene expression as we use the same starting material. More notably, this protocol could permit to perform ChIP/RNA-seq for kinetic experiment with short intervals (every minute or less) and to collect samples in the field.

Several studies have led to the identification of molecular mechanisms involved in dormancy, including a cluster of *DAM* genes (Bielenberg et al., 2008). *DAM*-related genes are up-regulated in dormant buds in peach (de la Fuente et al., 2015; Leida et al., 2012; Yamane et al., 2011), leafy spurge (Horvath et al., 2010), pear (Saito et al., 2015), apple (Mimida et al., 2015) and apricot (Sasaki et al., 2011). Conversely, *DAM*-related genes are down-regulated in non-dormant buds. In particular, it was shown that H3K27me3 abundance is increased at *DAM5* and *DAM6* loci in non-dormant buds compared to dormant buds (de la Fuente et al., 2015). Using the proposed ChIP-seq protocol, we found similar results for H3K27me3 abundance in the *DAM* genes in peach in non-dormant buds (Figure 2), thus validating our improved ChIP-seq method. In particular, we observe a higher enrichment for H3K27me3 at the *DAM5* and *DAM6* loci compared with *DAM3* in non-dormant buds in peach, suggesting that not all *DAM* genes are regulated the same way during dormancy. Additionally, we demonstrated the correlation between the presence of histone marks and gene expression in sweet cherry *(Prunus avium* L.) for two control genes *PavAG* and *PavEF1* known to be under control of H3K27me3 and H3K4me3, respectively (Figure 3c). As the ChIP-seq and RNA-seq were performed on the same biological material, the level of gene expression and the presence/absence of particular histone marks can be directly compared with confidence. In the last two decades ChIP has become the principal tool for investigating chromatin-related events at the molecular level such as transcriptional regulation. Our protocol will allow analysis of chromatin and expression dynamics in response to abiotic and biotic stresses, and this for trees in controlled conditions as well as growing in fields. Improvements to the ChIP-seq approach are still needed and will include an expansion of available ChIP-grade antibodies and a reduction of the hands-on time required for the entire procedure. A remaining challenge is to further decrease the amount of starting material without compromising the signal-to-noise ratio.

### Correlation between expression level and H3 K4 me 3 enrichment in sweet cherry dormant buds

Previous studies highlighted the importance of *DAM* genes as key components of dormancy in perennials. We have shown that, in sweet cherry, *PavDAM6* is highly expressed in October, at the beginning of dormancy, and then down-regulated over time (Figure 4). Conversely, we found that *PavDAM5* is highly expressed at the end of endodormancy (December), and then down-regulated during ecodormancy (Figure 4). Both *PavDAM5* and *PavDAM6* genes were down-regulated in ecodormancy, which suggest an important role in the maintenance of endodormancy as observed in Chinese cherry (Zhu et al., 2015), peach (Bielenberg et al., 2008; de la Fuente et al., 2015; Yamane et al., 2011) and Japanese apricot (Saito et al., 2015; Sasaki et al., 2011). However their timing of expression during endodormancy was different, suggesting that they have non-redundant roles and that their expression might be regulated by different factors during dormancy.

We conducted a ChIP-seq in sweet cherry for H3K4me3, which is associated with gene activation, in order to link the abundance of histone marks to expression patterns at three different dates along dormancy (October, December and January; Figure 4). We found an H3K4me3 enrichment around the translation start site of both *PavDAM6* and *PavDAM5* during dormancy (Figure 4), as reported in peach for *PpeDAM6* (*Leida et al., 2012*) (Leida et al. 2012) and in leafy spurge for *DAM1* (Horvath et al., 2010). Generally, H3K4me3 enrichment is more abundant at the *PavDAM5* locus than at the *PavDAM6* locus and this is associated with the difference in their expression levels (Figure 4). We find a positive relation between changes over time for H3K4me3 and expression level at *PavDAM5* (Figure 4). Similarly, we find a general trend that genes with an increase over time in H3K4me3 level also show an increase in expression, while genes with a reduction in H3K4me3 over time also show a reduction in expression (Figure 5). Despite these general correlations, changes in H3K4me3 signal that occur at a significant magnitude have been observed for only 671 genes. These results suggest that while chromatin marks might be involved in transcriptional regulation of some genes in sweet cherry buds during dormancy, other genes might exhibit changes in expression that are not associated with chromatin changes. In other genes, like *PavDAM6,* any relation between chromatin marks and expression seems to be more complex and possibly integrates other chromatin marks than H3K4me3. Performing a similar analysis for other chromatin marks, such as H3K27me3 that is associated with transcriptional repression, might reveal complex links between chromatin and expression dynamics during bud dormancy in sweet cherry.

Our analysis has been done with only 3 time-points spanning a period of four months. A finer time resolution would be needed to better identify relations between chromatin expression dynamics during dormancy, and in particular to define if chromatin changes happen before or after expression changes. Together with our results, the observation from Leida et al. (2012) and De la fuente et al. (2015) of the presence of repressive histone mark such as H3K37me3 in peach *DAM* loci during dormancy, support the hypothesis of a balance of histone mark enrichment controlling the dormancy process in sweet cherry and probably more largely in perennials.

### Genes involved in bud dormancy

In addition to *DAM* genes, we identified genes that showed differential H3K4me3 enrichment between the different bud dormancy stages. We found genes involved in the response to drought, cold and oxidative stresses that were either highly expressed at the beginning of endodormancy, such as *PavGSTU19* and *PavLEA14*, or expressed during ecodormancy (*PavXTH6*, *PavDAG2*, *PavGASA6* and *PavHVA22C)*. This is consistent with previous studies showing the key role of redox regulation (Ophir et al., 2009; Considine and Foyer, 2014; Beauvieux et al., 2018) and stress-associated stimulus (Maurya and Bhalerao 2017) in bud dormancy. Interestingly, we highlighted *PavGASA6* and *PavGA20ox1*, two genes associated with the gibberellin pathway. In particular, *PavGA20ox1*, that encodes an enzyme required for the biosynthesis of active GA (Plackett et al., 2012), is markedly upregulated after endodormancy release, associated with H3K4me3 enrichment at the locus, therefore suggesting that epigenetic and transcriptomic states facilitate an increase in active GA levels during ecodormancy, as previously shown in hybrid aspen (Rinne et al., 2011) and grapevine (Zheng et al., 2018).

Finally, our results highlighted the *CHROMATIN REMODELING5* (*PavCHR5*) gene, upregulated during ecodormancy. In Arabidopsis, *CHR5* is required to reduce nucleosome occupancy near the transcriptional start site of key seed maturation genes (Shen et al., 2015). We can therefore hypothesize that chromatin remodelling, other than post-transcriptional histone modification, occurs during dormancy progression. Our results confirm that the analysis of differential H3K4me3 states between endodormant and ecodormant buds might lead to the identification of key signalling pathways involved in the control of dormancy progression.

## MATERIALS AND METHODS

### Plant material

Sweet cherry trees (cultivar ‘Burlat’), apple trees (cultivar ‘Choupette’) and peach trees (unknown cultivar) were grown in an orchard located at the Fruit Experimental Unit of INRA in Toulenne (France, 44°34’N 0°16’W) under standard agricultural practices. Sweet cherry flower buds used for the RNA-seq and ChIP-seq experiment were collected on October 21st 2014 and December 5th 2014 for dormant buds and January 27th 2015 for non-dormant buds. Apple buds were collected on January 25th 2016 and peach buds on February 5th 2016. Buds were harvested from the same branches, flash frozen in liquid nitrogen and stored at −80°C prior to performing ChIP-seq and RNA-seq.

### Measurements of bud break

Three branches bearing floral buds were randomly chosen from the sweet cherry cultivar ‘Burlat’ trees at different dates. Branches were incubated in water pots placed in forcing conditions in a growth chamber (25°C, 16h light/ 8h dark, 60-70% humidity). The water was replaced every 3-4 days. After ten days under forcing conditions, the total number of flower buds that reached the BBCH stage 53 (Meier, 2001) was recorded. We estimate that endodormancy is released when the percentage of buds at BBCH stage 53 is above 50% after ten days under forcing conditions.

### ChIP/RNA-seq protocol

#### MATERIAL SAMPLING SECTION

Harvest tree buds in 2 ml tubes with screw cap, immediately flash-freeze in liquid nitrogen and store at −80°C until ready to proceed for the ChIP-seq or RNA-seq. There is no need to remove the scales after harvesting. Grind the tissues to a fine powder using mortars and pestles pre-chilled with liquid nitrogen. Add liquid nitrogen several times while grinding to facilitate cell lysis and to ensure that the material remains completely frozen to prevent degradation of tissues. Weights of powder for this protocol are for buds for which scales have not been removed.

For the cross-linking and chromatin extraction, weigh out 300 to 500 mg of powder in a 50 ml Falcon tube pre-chilled with liquid nitrogen. The same amount of powder should be used for all samples to allow a direct comparison of results. Then proceed to “ChIP and library preparation section”.

For RNA extraction, weigh out 50-70 mg of powder in a 2 ml tubes (screw cap) pre-chilled with liquid nitrogen. Then proceed to “RNA extraction and library preparation section”. Due to the small amount of starting material, it is necessary to keep the tubes in liquid nitrogen to prevent any degradation.

#### ChIP AND LIBRARY PREPARATION SECTION

**Cross-linking and chromatin extraction:** Timing 2-3 hours. Work on ice, except when specified otherwise.

Add 25 ml of ice-cold Extraction buffer 1 [0.4 M sucrose, 10 mM HEPES pH 7.5, 10 mM MgCl_2_, 5 mM β-mercaptoethanol, 1 mM PMSF, 1 % PVP-40 (polyvinylpyrrolidone), 1 tablet of complete protease inhibitor EDTA free for 50 ml of buffer from Sigma cat# 11836170001] to the powder. For each buffer, the protease inhibitor (PMSF), the tablet of complete protease inhibitor and β-mercaptoethanol should be added directly before using the buffer. PVP-40 is a chelator used to remove phenolic derivatives as well as polysaccharides and improve the quality of the chromatin extraction. It is commonly used in RNA and chromatin extractions in buds sampled from fruit trees (Gambino et al., 2008; Leida et al., 2012; Ionescu et al., 2017). When extracting chromatin in other biological systems such as *Arabidopsis thaliana* or *Saccharomyces cerevisiae*, the PVP-40 in Extraction buffer 1 may optionally be removed.

Immediately add 675 µl of 37% formaldehyde solution (1% final concentration) and invert the tube several times to resuspend the powder. Cross-link the samples by incubating at room temperature for 10 minutes and then quench the formaldehyde by adding 1.926 ml 2 M of fresh glycine solution (0.15 M final concentration). Invert the tube several times and incubate at room temperature for 5 minutes. Filter the homogenate through Miracloth (Millipore cat# 475855) in a funnel and collect in a clean 50 ml Falcon tube placed on ice. Repeat the filtration step once more. Centrifuge the filtrate at 3,200 × g for 20 minutes at 4 °C. Discard the supernatant by inverting the tube, being careful not to disturb the pellet. Gently resuspend the pellet in 1 ml of Extraction buffer 2 [0.24 M sucrose, 10 mM HEPES pH 7.5, 10 mM MgCl_2_, 1 % Triton X-100, 5 mM β-mercaptoethanol, 0.1 mM PMSF, 1 tablet protease inhibitor EDTA free for 50 ml of solution], without creating bubbles, and transfer the solution to a clean 1.5 ml tube. Centrifuge at 13,500 × g for 10 minutes at 4°C. Carefully remove the supernatant by pipetting. If the pellet is still green, repeat the resuspension in 1 ml of Extraction buffer 2, centrifuge at 13,500 × g for 10 minutes at 4°C, and remove the supernatant. In a new 1.5 ml tube, add 300 µl of Extraction buffer 3 [1.7 M sucrose, 10 mM HEPES pH 7.5, 2 mM MgCl_2_, 0.15 % Triton X-100, 5 mM β-mercaptoethanol, 0.1 mM PMSF, 1 mini-tablet protease inhibitor EDTA free for 50 ml of solution]. Slowly resuspend the pellet in 300 µl of Extraction buffer 3 to prevent the formation of bubbles. Take the resuspended pellet and carefully layer it on top of the 300 µl Extraction buffer 3. Centrifuge at 21,200 × g for 1 hour at 4°C. During this process, nuclei are pelleted through a sucrose cushion to remove cellular contaminants.

From this step, chromatin fragmentation can be performed in two different ways: (A) sonication, to shear the chromatin into 100-500 bp fragments, or (B) MNase (Micrococcal nuclease) digestion, to enrich for mono-nucleosomes (∼150-200 bp). Results shown in this paper come from ChIP-seq performed on sonicated chromatin.

##### A. Sonication: TIMING 3-4h hours (+ 8 hours of incubation)

###### i. Chromatin fragmentation

Carefully remove the supernatant with a pipette and resuspend the nuclei pellet in 300 µl of Sonication buffer [50 mM HEPES pH 7.5, 10 mM EDTA, 1 % SDS, 0.1 % sodium deoxycholate, 1% Triton X-100, 1 mini-tablet protease inhibitor EDTA free for 50 ml of solution]. To improve nuclear membrane breaking, flash-freeze the tube in liquid nitrogen and then thaw rapidly by warming the tube in your hand. Repeat once more. This freeze-thaw step is not essential, but could improve the chromatin yield. Centrifuge the tube at 15,800 × g for 3 minutes at 4°C to pellet debris, and carefully recover the supernatant into a new tube. Complete the tube to 300 µl with the Sonication buffer. Set aside a 10 µl aliquot of chromatin in a PCR tube to serve as the non-sonicated control when assessing sonication efficiency by gel electrophoresis and keep on ice. Shear the chromatin into ∼300 bp (100–500 bp) fragments by sonication (e.g. using Diagenode Bioruptor Twin-UCD400, sonicate 300 µl chromatin in 1.5 ml microcentrifuge tubes for 14 to 16 cycles, on High setting, with 30s ON/30s OFF per cycle). The number of cycles of sonication to obtain DNA fragments of around 300 bp should be tested and optimised for different tissues and different concentrations of chromatin. Transfer 40 µl of sheared chromatin to a PCR tube, which will be used to check the sonication efficiency. The rest of the sonicated chromatin should be stored at −80°C.

###### ii. Analysis of sonication efficiency

Complete the sonicated (40 µl) and non-sonicated (10 µl) aliquots to 55.5 µl with TE buffer [10 mM Tris-HCl pH 8, 1 mM EDTA], add 4.5 µl of 5 M NaCl and incubate at 65°C for 8 hours to reverse cross-link. Add 2 µl of 10 mg/ml RNase A (Fisher cat# EN0531) and incubate at 37°C for 30 minutes. Add 2 µl of 20 mg/ml proteinase K (Fisher cat# EO0491) and incubate at 45°C for 1 hour. During this step, take out the SPRI beads (e.g AMPure beads; Beckman Coulter, cat# A63880) from the fridge and allow them to equilibrate at room temperature (for at least 30 minutes before use).

To extract DNA using SPRI beads, vortex the beads until they are well dispersed, add 126 µl of beads to 60 µl of sample (2.1 × ratio) and mix well by pipetting up and down at least 10 times. Incubate 4 minutes at room temperature and then place the tubes on a magnetic rack (96 well; Fisher, cat# AM10027) for 4 minutes to capture the beads. Carefully remove and discard the supernatant without disturbing the beads. Without removing the tubes of the magnetic rack, add 200 µl of freshly prepared 80 % v/v ethanol, incubate for 30 seconds and discard the supernatant. Repeat the ethanol wash once more and then completely remove all ethanol. Allow the beads to dry for 15-30 minutes, until cracks appear in the bead pellet and no droplets of ethanol are visible. Tubes can alternatively be placed in a fume hood for 10 minutes to accelerate drying. The beads must be completely free from ethanol as it can interfere with downstream processes. Remove the tubes from the magnetic rack and resuspend the beads in 15 µl of 10 mM Tris-HCl (pH 8) by pipetting up and down at least 10 times. Incubate for 5 minutes at room temperature and place on the magnetic rack for 4 minutes to capture the beads. Carefully transfer 14 µl of supernatant containing DNA to a new tube. Add 2.8 µl of 6× Loading dye to 14 µl of DNA. Separate the DNA by electrophoresis on a 1.5 % agarose gel for at least 1 h at 70 V. The smear should be concentrated between 100–500 bp (Figure 6a). If necessary, perform additional sonication cycles. Otherwise, continue directly to the “Immunoprecipitation (IP)” step.

**Figure 6.**
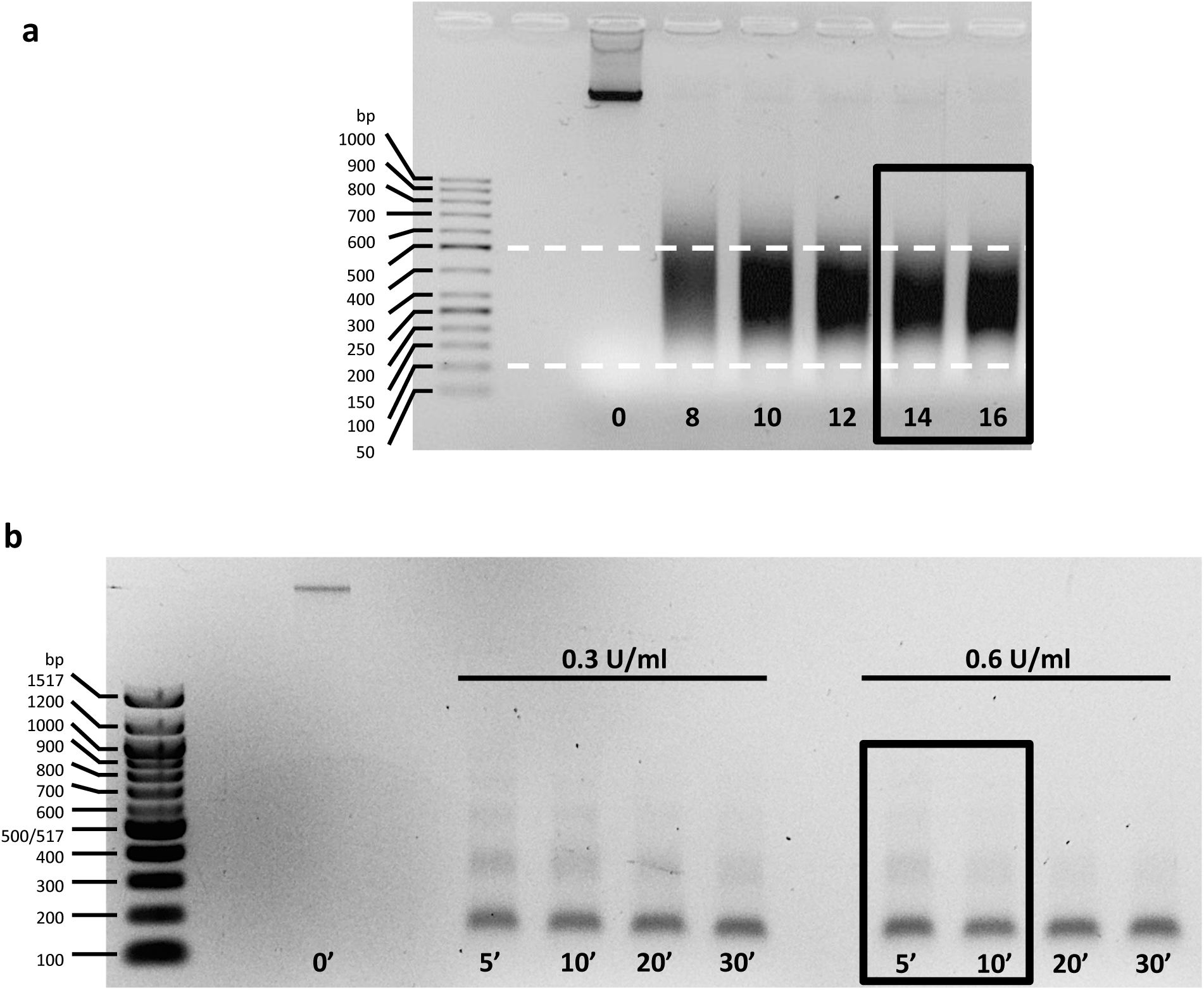
DNA profiles for MNase-digested and sonicated chromatin (*Prunus avium* L.) a) DNA profiles of chromatin fragmented with different numbers of sonication cycles (0-16). The dotted white lines represent the optimal range of DNA fragments size (100-500 bp). Optimal sonication profiles are indicated with the black rectangle. b) DNA profiles of chromatin digested with different concentrations of MNase (0.3 and 0.6 U/ml) for various durations (5, 10, 20 and 30 minutes). Optimal MNase digestion profiles, with ∼80% of chromatin in mono-nucleosome form, are indicated with the black rectangle.

##### B. MNase digestion: TIMING 4-5 hours (+ 2 *×* 8 hours of incubation)

###### i. DNA quantification prior to MNase digestion

Carefully remove the supernatant with a pipette and resuspend the nuclei pellet in 500 µl of MNase buffer [20 mM HEPES pH 7.5, 50 mM NaCl, 0.5 mM DTT, 0.5 % NP-40, 3mM CaCl_2_, Triton X-100, 1 mini-tablet protease inhibitor EDTA free for 50 ml of solution]. To improve nuclear membrane breaking, flash-freeze the tube in liquid nitrogen and then thaw rapidly by warming the tube in your hand. Repeat once more. This freeze-thaw step is not essential, but could improve the chromatin yield. Transfer 40 µl of chromatin to a PCR tube to quantify DNA prior to MNase digestion and complete to 55.5 µl with MNase digestion buffer, add 4.5 µl of 5M NaCl and incubate in a PCR machine or thermocycler at 65°C for 8 hours to reverse cross-link. Keep the rest of the chromatin at –80 °C. Add 2 µl of 10 mg/ml RNase A and incubate at 37°C for 30 minutes. Add 2 µl of 20 mg/ml proteinase K and incubate at 45°C for 1 hour. During this step, take out the SPRI beads from the fridge and allow them to equilibrate at room temperature (for at least 30 minutes before use). Proceed to the DNA extraction using SPRI beads as explained before in the sonication analysis section (ii). Use 1 µl from each sample to quantify the DNA using a Qubit fluorometer (ThermoFisher Scientific), or a Nanodrop spectrophotometer (Thermo Scientific).

###### ii. MNase digestion

Adjust all samples to the same concentration according to the quantification results using MNase buffer and to a final volume of 500 µl. Set aside a 20 µl aliquot of chromatin in a PCR tube to serve as the non-digested control when assessing MNase efficiency by gel electrophoresis and keep on ice. Incubate chromatin in a ThermoMixer (Eppendorf) for 2-5 minutes at 37°C with shaking at 1,200 rpm, for optimal MNase activity. Add MNase (Fisher cat# 88216) to the chromatin to a final concentration of 0.6 U/ml and incubate 10 minutes in the ThermoMixer at 37°|C, 1,200 rpm. Stop the digestion by adding 5 µl of 0.5 M EDTA pH 8 (5 mM final concentration), invert the tube several times to mix and immediately place on ice for 5 minutes. The optimal MNase enzyme concentration and incubation time to obtain predominantly mono-nucleosomes should be tested and optimised for different tissues and different concentrations of chromatin. For the optimisation of MNase digestions, we recommend using 1 ml of chromatin and carrying out digestions in 100 µl aliquots with varying concentrations of MNase (0.2 U/ml to 1 U/ml) and incubation times (5 to 20 minutes). Transfer 50 µl of digested chromatin to a PCR tube, which will be used to check the MNase efficiency. The rest of the digested chromatin should be stored at −80°C.

###### iii. MNase digestion analysis

Complete the digested sample (50 µl) and non-digested (20 µl) aliquots to 55.5 µl with TE buffer, add 4.5 µl of 5M NaCl and incubate in a PCR machine or thermocycler at 65°C for 8 hours to reverse cross-link. Add 2 µl of 10 mg/ml RNase A and incubate at 37°C for 30 minutes. Add 2 µl of 20 mg/ml proteinase K and incubate at 45°C for 1 hour. During this step, take out the SPRI beads from the fridge and allow them to equilibrate at room temperature (for at least 30 minutes before use). Proceed to the DNA extraction, using SPRI beads as explained before in the sonication analysis section (ii). Add 2.8 µl of 6× Loading dye to 14 µl of DNA. Separate the DNA by electrophoresis on a 1.5% agarose gel for at least 1 h at 70 V. The most abundant band should be 150-200 bp in size (Figure 6b), which corresponds to chromatin in mono-nucleosome form, with a less abundant 300-350 bp band (di-nucleosomes) and a faintly visible ∼500 bp band (tri-nucleosomes). For optimum sequencing results, approximately 80% of chromatin should be in mono-nucleosome form. If this is judged not to be the case from the gel, it is not possible to carry out further MNase digestions on the chromatin, as EDTA sequesters calcium ions that are required for MNase activity.

**Immunoprecipitation (IP):** TIMING 7-8 hours (+ overnight incubation). Work on ice, except when specified otherwise.

Transfer 50 µl of protein A- and/or protein G-coupled magnetic beads (Invitrogen cat# 10-002D and cat# 10-004D, respectively) per IP to a 2 ml tube. Wash the beads with 1 ml of Binding buffer [0.5% (wt/vol) BSA, 0.5% (vol/vol) Tween-20 in PBS (without Ca_2_^+^, Mg_2_^+^)] during 5 minutes at 4°C on a rotating wheel (low speed, around one rotation every 5-6 seconds). Place the tubes on a magnetic rack (Thermo Fisher cat# 12321D) until the liquid is clear and remove the supernatant. Repeat three times. After the washes, resuspend the beads in 250 µl of Binding buffer. Add 5 µl of antibody per IP to the beads. In our study, we used anti-trimethyl-histone 3 Lys 27 antibody (Millipore cat# 07-449) and anti-trimethyl-histone 3 Lys 4 antibody (Millipore cat#17-614,). Incubate 4 hours on a rotating wheel at 4°C (low speed). During this incubation time, centrifuge the sonicated or digested chromatin at 15,800 × g for 5 minutes at 4°C to pellet debris, and carefully recover supernatant into a new tube. Transfer 100 µl of sonicated chromatin to a new 2 ml tube for one IP, add 900 µl of Binding buffer and keep on ice. Or transfer 200 µl of MNAse-digested chromatin to a new 2 ml tube for one IP, add 800 µl of Binding buffer and keep on ice. Transfer 20 µl of sonicated or MNAse-digested chromatin as an input fraction (no immunoprecipitation) in a PCR tube and store at −80°C. The rest of the chromatin can be used for another IP or stored at −80°C. After completion of the incubation of protein A/G beads with the antibody, place the tubes containing the antibody-bead complexes on a magnetic rack, remove the supernatant and wash the beads with 1 ml of Binding buffer during 5 minutes at 4°C on a rotating wheel (low speed). Repeat three times. Resuspend the beads in 50 µl of Binding buffer per IP and transfer to the 1 ml of diluted chromatin. Incubate overnight on a rotating wheel at 4°C (low speed). Briefly centrifuge the tube (<3 seconds) to pull down the liquid in the lid of the tube. Place on a magnetic rack and remove the supernatant. Wash the beads to reduce unspecific interactions by incubating 5 minutes at 4°C on a rotating wheel (low speed) with 1ml of the following buffers and total number of washes:

a. 5 washes with Low Salt Wash buffer [150 mM NaCl, 0.1% SDS, 1% triton X-100, 2 mM EDTA, 20 mM Tris-HCl pH 8];
b. 2 washes with High Salt Wash buffer [500mM NaCl, 0.1% SDS, 1% triton X-100, 2 mM EDTA, 20mM Tris-HCl pH 8];
c. 2 washes with LiCl Wash buffer [0.25 M LiCL 1% NP-40 (IGEPAL), 1% sodium deoxycholate, 1 mM EDTA, 10 mM Tris-HCl pH 8];
d. 2 washes with TE buffer.

After the second wash in TE buffer, resuspend the beads in 100 µl of TE buffer and transfer the beads to a PCR tube. Place the tube on a magnetic rack, remove the TE buffer and resuspend the beads in 60 µl of Elution buffer [10 mM Tris-HCl pH 8.0, 5 mM EDTA pH 8.0, 300 mM NaCl, 0.5% SDS].

**Reverse cross-linking and Elution by proteinase K treatment:** TIMING 2 hours (+ 8 hours of incubation)

Defrost the input fraction on ice (20 µl of sonicated or MNAse-digested chromatin). Complete the input fraction to 60 µl with the Elution buffer. Incubate the input fraction and the IP sample (60 µl beads-Elution buffer) at 65°C for 8 hours in a PCR machine or thermocycler to reverse crosslink. Add 2 µl of RNase A (10 mg/ml) and incubate at 37°C for 30 minutes. Add 2 µl of Proteinase K (20 mg/ml) and incubate at 45°C for 1 hour. During this step, take out the SPRI beads from the fridge and allow them to equilibrate at room temperature (for at least 30 minutes before use). Place the tubes on a magnetic rack to collect the beads, transfer 60 µl of supernatant from each well to a new PCR tubes (or a new 96 wells-plate).

**DNA extraction using SPRI beads and q PCR:** TIMING 1 hour

Proceed to the DNA extraction, using SPRI beads as explained before in the sonication analysis section (ii) until just before the elution. Remove the tubes from the magnetic rack and for the elution, resuspend the beads in 50 µl of 10mM Tris-HCl (pH 8.0) by pipetting up and down at least 10 times. Incubate for 5 minutes at room temperature and place on the magnetic rack for 4 minutes to capture the beads. Carefully transfer 49 µl of supernatant containing DNA to a new tube. For qPCR analysis, use 1 µl of DNA per 10 µl reaction, from the IP and input. The percentage of enrichment of DNA in the ChIP fraction relative to the input fraction is calculated according to the formula: (2^-Cp ChIP^ / 2 ^-Cp input^) × 100. Keep the rest of the DNA for sequencing (continue to “ChIP library preparation and size selection section”).

**ChIP library preparation and size selection:** TIMING 2-3 days

Use 5–10 ng from the input fraction for the preparation of sequencing libraries. Quantify the input fraction using a DNA high sensitivity Qubit kit. For the IP fraction, as yield is often too low to be able to quantify DNA, we recommend to use the entire volume from the IP for the preparation of sequencing libraries. Using this protocol we extracted around 500 ng of DNA from 300 to 500 mg of powder of sweet cherry buds. We recommend carrying out ChIP-seq library preparation using the TruSeq ChIP Sample Prep Kit (Illumina 48 samples, 12 indexes, Illumina, cat# IP-202-1012) with minor modifications.

1. The “Purify Ligation Products” section using the gel electrophoresis is eliminated to minimise DNA loss.
2. DNA size selection is carried out after the “Enrich DNA fragments” section. This is a required step to increase the visualisation of nucleosome positioning. Smaller and larger reads might disturb the MNase input profile after analysis.

The Illumina TruSeq ChIP Sample Preparation protocol is available at the following URL: http://support.illumina.com/content/dam/illumina-support/documents/documentation/chemistry_documentation/samplepreps_truseq/truseqchip/truseq-chip-sample-prep-guide-15023092-b.pdf

Check the quality of libraries using a 4200 TapeStation or Bioanalyzer instruments (Agilent) following manufacturer’s. See Suppl. Figure 2 for profiles with and without adapter contaminations. If the libraries are contaminated with adapters, repeat again a size selection step with SPRI beads to remove them, otherwise proceed directly with the “quantification and pool of libraries” section. Store DNA for sequencing at −20°C.

#### RNA EXTRACTION AND LIBRARY PREPARATION SECTION

**RNA extraction:** TIMING 2-5 hours

We recommend for the RNA extraction the use of RNeasy® Plant Mini kit from Qiagen (cat# 74904) for less than 50 samples with the following minor modifications:

1. Start from 50-70 mg of buds powder with scales. Only remove the tubes from the liquid nitrogen when the RNA Extraction buffer is prepared.
2. Add 1.5 % of PVP-40 (polyvinylpyrrolidone) in the RLT buffer to chelate phenolic compounds and thus prevent any interaction. Then add the appropriate volume of ß-mercaptoethanol mentioned in the Qiagen protocol.
3. Add 750 µl of RNA Extraction buffer (RLT buffer + PVP-40 + ß-mercaptoethanol) instead of 450 µl if the starting material contains scales to increase the RNA yield.

Alternatively, RNA can be extracted using the MagMAX™-96 Total RNA Isolation Kit from Thermo Fisher (cat# AM1830) for more than 50 samples following manufacturer’s instructions. Store RNA at −80°C.

**RNA library preparation:** TIMING 3-4 days

We recommend carrying out RNA-seq library preparation using the Truseq Stranded mRNA Library Prep Kit from Illumina (96 samples, 96 indexes, Illumina cat# RS-122-2103). Check the quality of libraries using 4200 TapeStation or Bioanalyzer instruments (Agilent) following manufacturer’s instructions. See Suppl. Figure 3 for profiles of libraries with and without adapter contaminations. If the libraries are contaminated with adapters, continue with a size selection step with SPRI beads to remove them, otherwise proceed directly with “the quantification and pool of libraries” section.

#### QUANTIFICATION AND POOL OF LIBRARIES SECTION

**Quantification of RNA and ChIP libraries:** From this step, the quantification and the pool for RNA and ChIP libraries are the same. However, ChIP-seq libraries on one-hand and RNA-seq libraries on the other should be pooled and sequenced separately. Libraries are quantified using a Qubit Fluorometer from Thermo Fisher (DNA high sensitivity). Dilute the DNA high sensitivity dye to 1/200 in the DNA high sensitivity buffer (e.g. for 10 samples: mix 1.990 ml of DNA high sensitivity buffer and 10 µl of DNA high sensitivity dye). Add 198 µl of mix in Qubit tubes (Thermo Fisher cat# Q32856) and 2 µl of DNA (for standard: 190 µl of mix + 10 µl of standard). Vortex and spin down. Quantification is performed using Qubit fluorometer following manufacturer’s instructions.

**Pool of libraries:** TIMING 1 hour

According to the quantification results, dilute libraries at 10 nM using this calcul to convert from ng/µl to nM: (concentration*10^6)/(size*617.96+36.04), where concentration is in ng/µl and size in bp. And pool the libraries using 5 µl of each library. Quantify the pool by Qubit as explained before and dilute the pool to the concentration required by the sequencing facility/company or the sequencer system used.

### Data analysis

#### (i) RNA-seq

The raw reads obtained from the sequencing were analysed using several publicly available software and in-house scripts. Firstly, we determined the quality of reads using FastQC (www.bioinformatics.babraham.ac.uk/projects/fastqc/). Then, possible adaptor contaminations were removed using Trimmomatic (Bolger et al., 2014), before alignment to the *Prunus persica* v.1 (Verde et al., 2017) or *Malus domestica* v.3 (Velasco et al., 2010) reference genome using Tophat (Trapnell et al., 2009). Possible optical duplicates resulting from library preparation were removed using the Picard tools (https://github.com/broadinstitute/picard). Raw reads and TPM (Transcripts Per Million) were computed for each gene (Wagner et al., 2012). To finish, data are represented using the Integrative Genome Viewer (Robinson et al., 2011) as a tool for visualising sequencing read profiles.

#### (ii) ChIP-seq

Sequenced ChIP-seq data were analysed in house, following the same quality control and pre-processing as in RNA-seq. The adaptor-trimmed reads were mapped to the *Prunus persica* reference genome v.1 (Verde et al., 2017) or *Malus x domestica* reference genome v3.0 (Velasco et al., 2010) using Bowtie2 (Langmead et al., 2009). Possible optical duplicates were removed using Picard, as described above. Data are represented using the Integrative Genome Viewer (Robinson et al., 2011). The efficiency of the H3K4me3 ChIP-seq was evaluated using fingerplots (Suppl. Figure 6a) from deeptools (Ramirez et al., 2016). We observe that a smaller fraction of the genome contains a high proportion of reads for all H3K4me3 ChIP-seq compared to H3 ChIP-seq, suggesting that the H3K4me3 ChIP worked. Also, around 30% of the genome is not covered by reads, which can be explained by the fact that the ChIP-seq has been performed on cherry tree buds, but mapped on the peach genome.

We used a gene-centric approach to identify genes with significant changes in the strength of H3K4me3 signal between time-points. The DiffBind R package was used for the read counting and differential binding analysis steps (Stark and Brown, 2011; Ross-Innes et al., 2012). Firstly, to quantify H3K4me3 signal at genes, we measured the number of H3K4me3 ChIP reads in a 2000bp window around the TSS of each gene (Suppl. Figure 7a). Two biological replicates were present for each time point, and DiffBind combines the available information to provide a statistical estimate of the H3K4me4 signal for any particular gene. Next, we identified a subset of genes that exhibit significant differential binding between any two time-points (October vs December; October vs January and December vs January). For this step, we used the H3 ChIP as a control, instead of the INPUT, because the number of mapped reads for one of the INPUT is much lower than other samples (Suppl. Table 1) and this might have created some bias in detecting genes with significant differences in H3K4me3 enrichment between time-points. The quality of biological replicates was assessed by performing a correlation heatmap, and hierarchical clustering of samples (Suppl. Figure 7b), based on the H3K4me3 signal around TSS for all genes, normalised by H3. It shows that H3K4me3 ChIP-seq replicates are of high quality.

To identify groups of genes with similar H3K4me3 dynamics, hierarchical clustering was performed on the Z-score of the H3K4me3 signal normalised by H3 using the function hclust on 1-Pearson correlation in the statistical programme R (R Core Team 2014). The Z-score has the formula (signal for a time-point - average over all time-points/ STD across all time-points), which allows the changes in H3K4me3 between time-points to be compared on a gene-to-gene basis, after normalising for differences that exist between genes.

## Declarations

### Data Archiving Statement

ChIP-seq and RNA-seq raw data will be made available on GEO upon acceptation of the manuscript.

### Authors’ contributions

SC and BW organized the project. NV performed the experiments and analysed the data. DGS performed the ChIP-qPCR in *Saccharomyces cerevisiae*, and SC performed the ChIP-qPCR in *Arabidopsis thaliana*. NV, SC and BW wrote the paper. DGS, FR, ED and PAW edited the paper. All authors read and approved the final manuscript.

## Acknowledgments

We thank the Fruit Experimental Unit of INRA (Bordeaux-France) for growing the trees and Varodom Charoensawan (Mahidol University, Thailand) for sharing his scripts for RNA-seq and ChIP-seq mapping.

---

**Supplemental figure 1.**
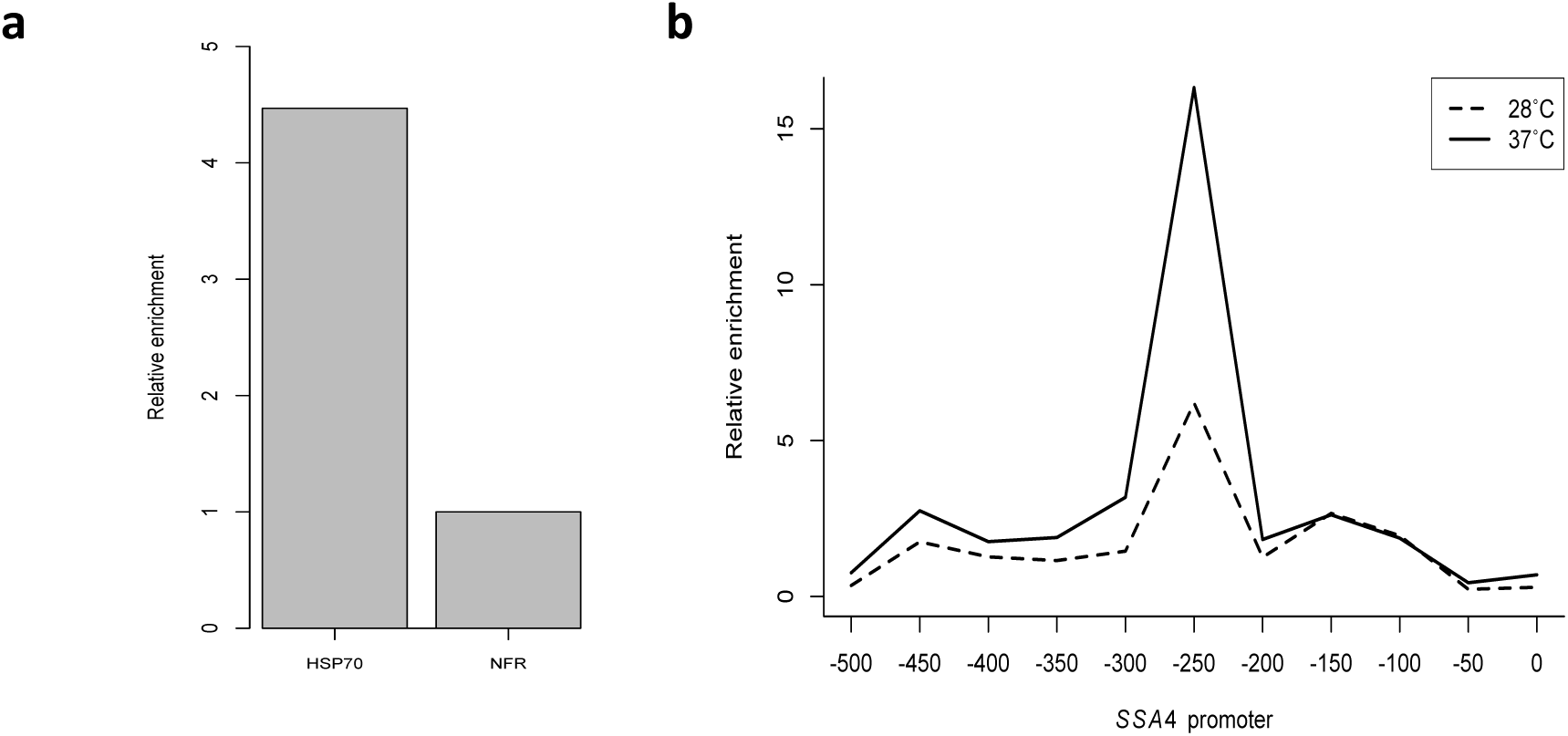
ChIP results in different biological systems. a) ChIP-qPCR on *Arabidopsis thaliana* seedlings. 7-day-old Col-0 seedlings grown at 22°C were harvested after 15-minute incubation at 17°C and flash frozen. ChIP was performed as outlined in the protocol, with an anti-HTA9 antibody (44). Quantitative PCR was carried out using primers specifically amplifying the genomic region occupied by the +1 nucleosome of the HSP70/ AT3G12580 or the upstream nucleosome-free region (NFR). Occupancy is normalised to the input fraction. b) ChIP-qPCR on *Saccharomyces cerevisiae* (budding yeast). YEF473a cells expressing FLAG×3-tagged Hsf1 were grown in YPD medium untill mid log phase at 28°C and subjected to 5-minute heat treatment at 37°C, after which they were harvested by centrifugation and flash frozen. ChIP was performed as outlined in the protocol, with an anti-FLAG antibody. Quantitative PCR was carried out using primers specifically amplifying distinct regions of the promoter of Hsf1-target gene SSA4 (denoted as distance from the start of the coding sequence). Occupancy is normalised to the input fraction.

**Supplemental figure 2.**
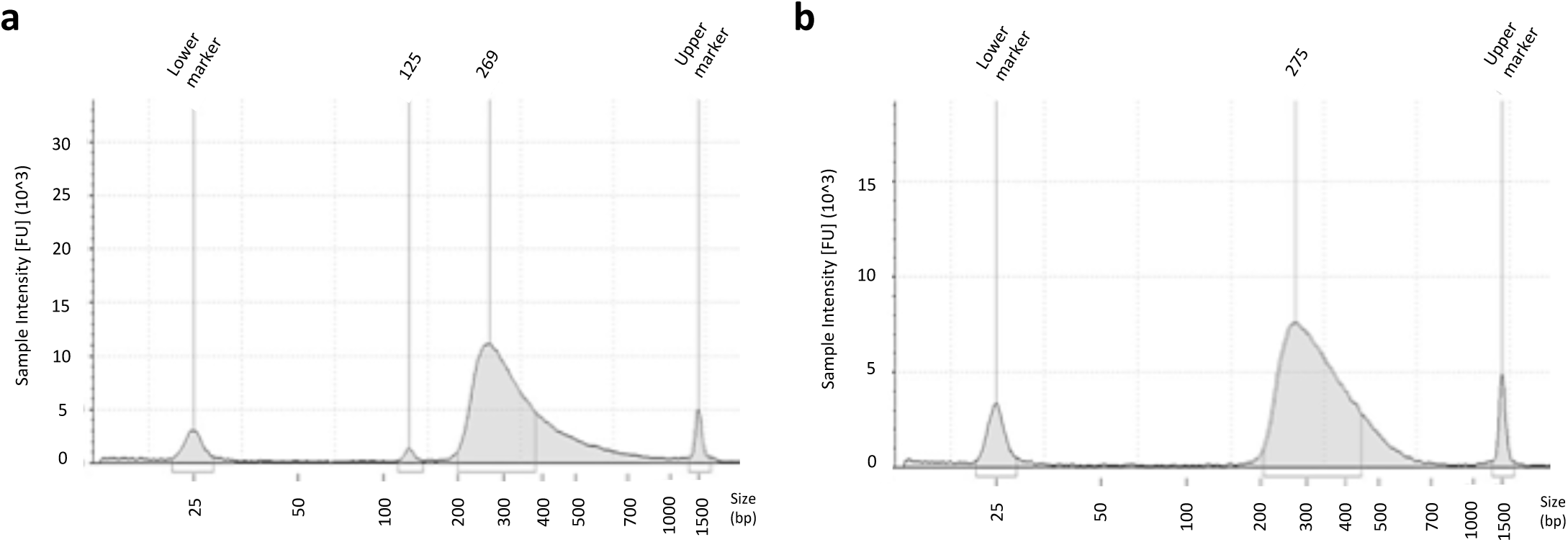
Example of ChIP-seq library profile with (a) and without (b) adapter contamination. The peak at 125 bp corresponds to adapter contaminations. A size selection using SPRI beads is carried out to remove the adapter contamination prior sequencing. DNA profiles were obtained using TapeStation 4200 (Agilent Genomics).

**Supplemental figure 3.**
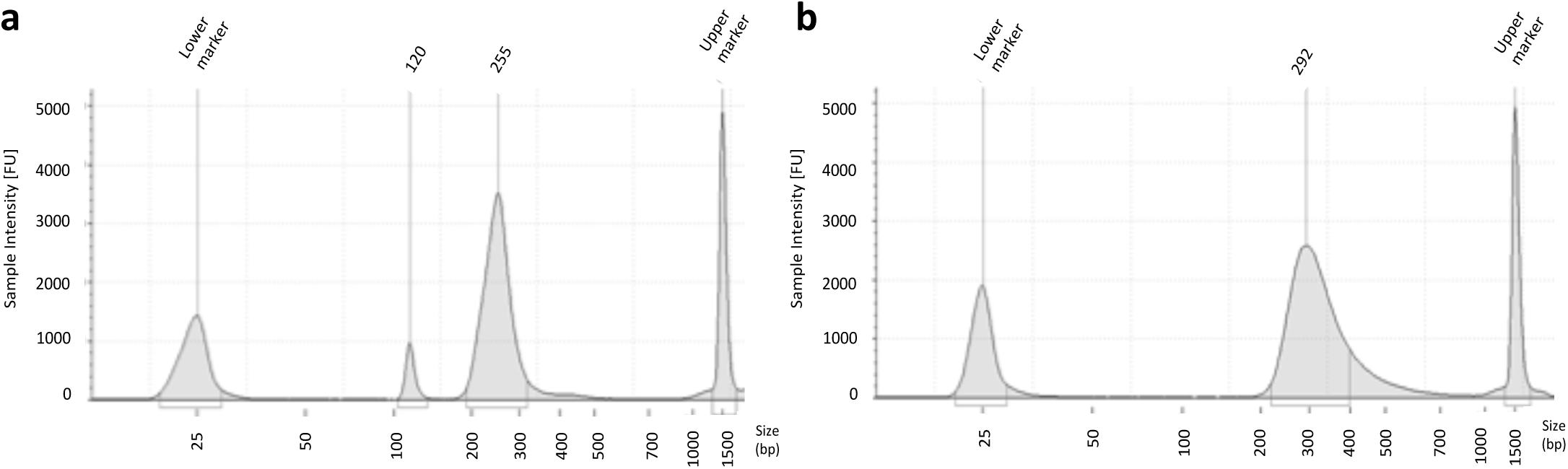
Example of RNA-seq library profile with (a) and without (b) adapter contamination. The peak at 120 bp corresponds to adapter contaminations. A size selection using SPRI beads is carried out to remove the adapter contamination prior sequencing. DNA profiles were obtained using TapeStation 4200 (Agilent Genomics).

**Supplemental figure 4.**
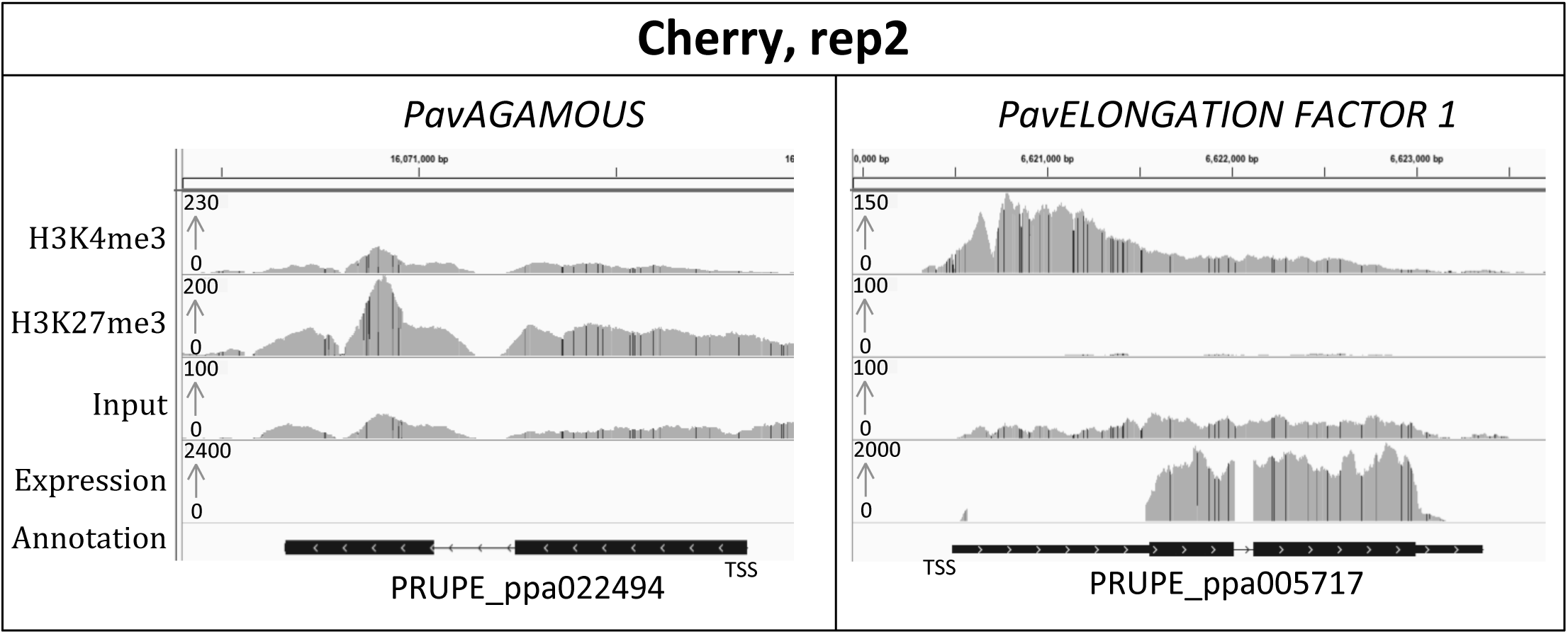
IGV screenshot of ChIP-seq data for H3K27me3 and H3K4me3 their corresponding input and expression in sweet cherry buds. Replicate 1 is shown in Figure 3c. Genes are represented by black rectangles, with arrows indicating gene directionality and taller boxes within the rectangles representing exons.

**Supplemental figure 5.**
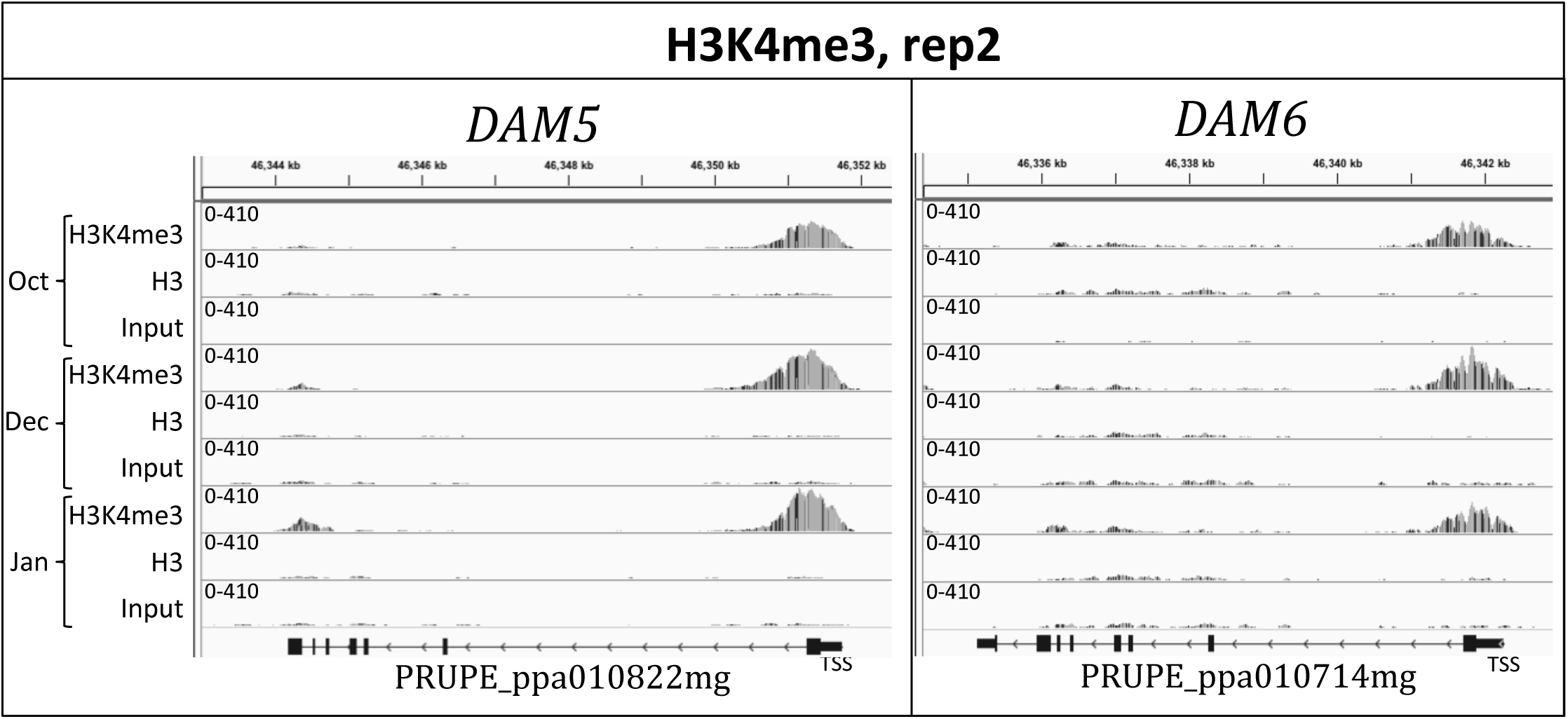
IGV screenshot of the second biological replicate of ChIP-seq data for H3K4me3 and H3 at three different dates (21st October 2014, 5th December 2014 and 27th January 2015) and their corresponding inputs at *PavDAM5* and *PavDAM6* loci. Replicate 1 is shown in Figure 4c. Genes are represented by black rectangles, with arrows indicating gene directionality and taller boxes within the rectangles representing exons.

**Supplemental figure 6.**
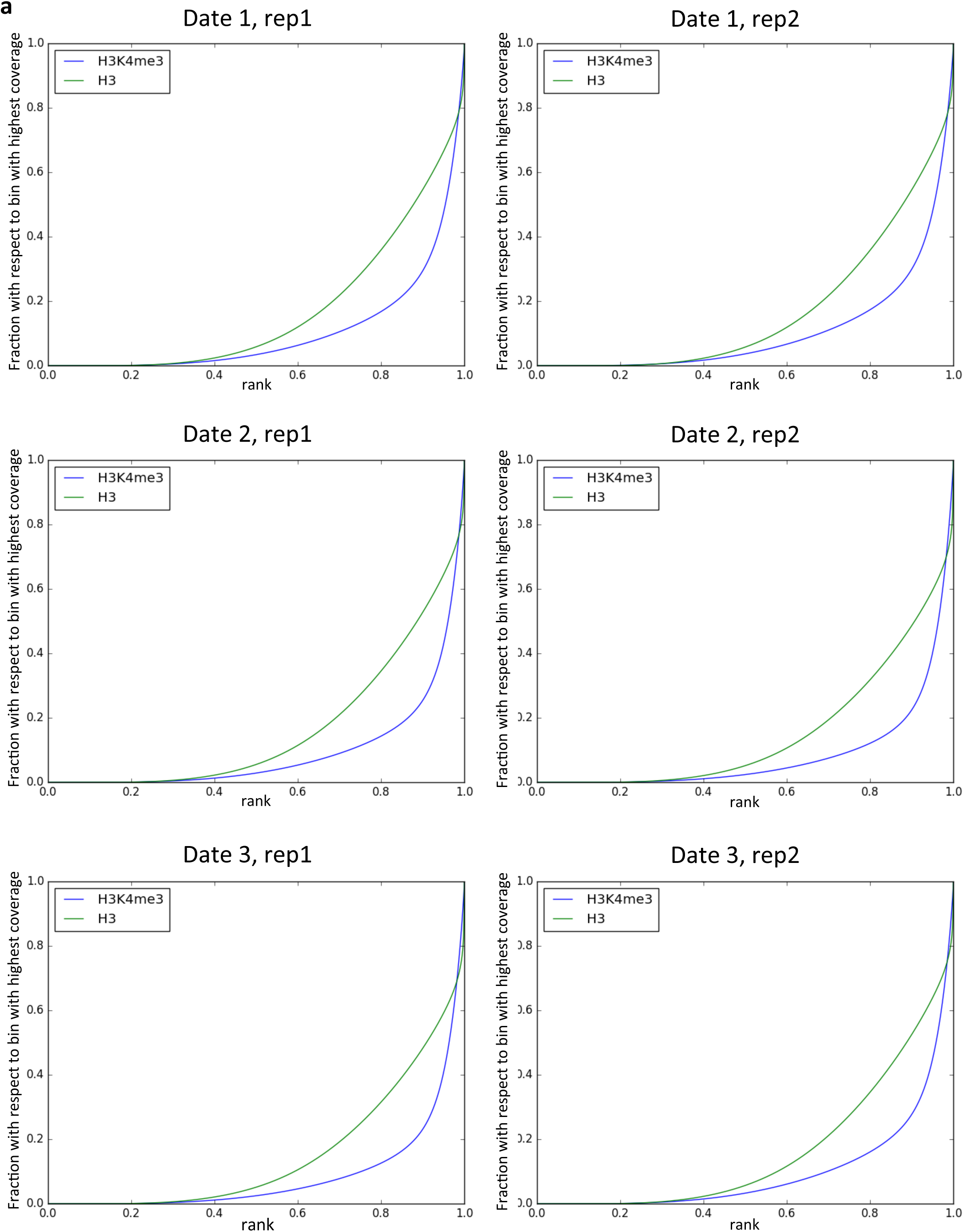
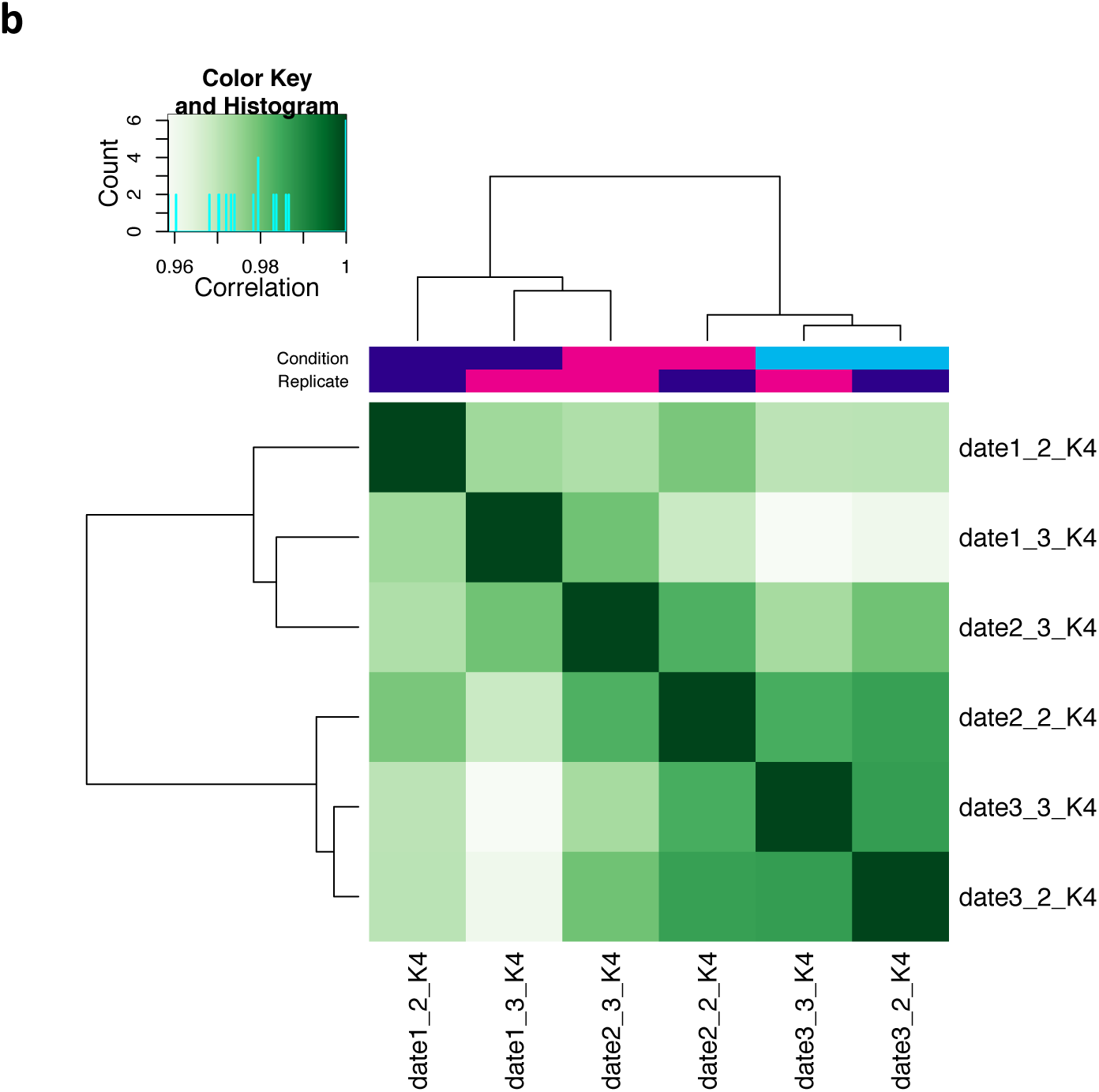
a) Fingerplots of H3 and H3K4me3 ChIP-seq. Each plot is for a replicate of a time-point. b) Heatmap of correlation between H3K4me3 ChIP-seq samples. The correlation is based on the H3K4me3 signal around TSS for all genes, normalised by H3.

**Supplemental figure 7.**
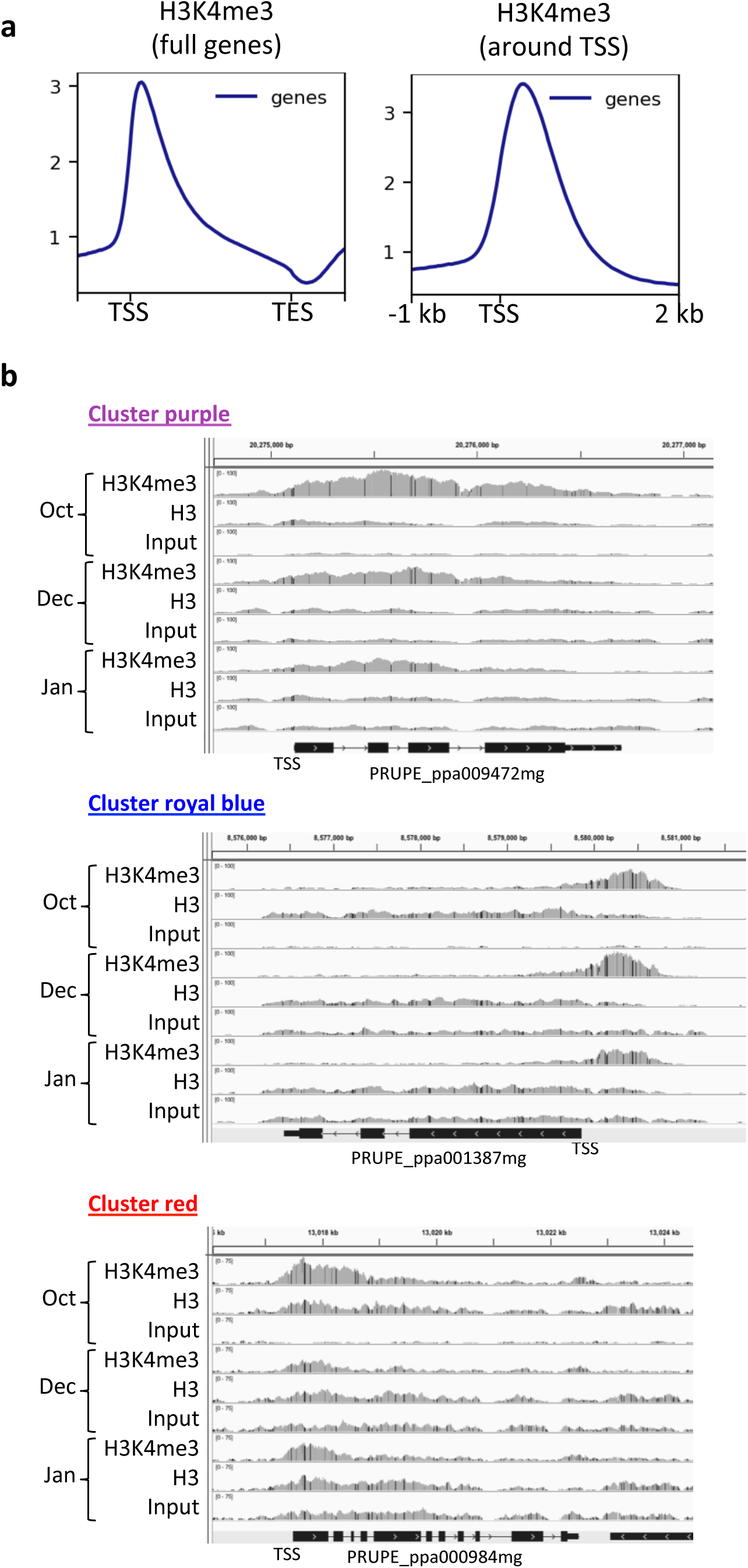
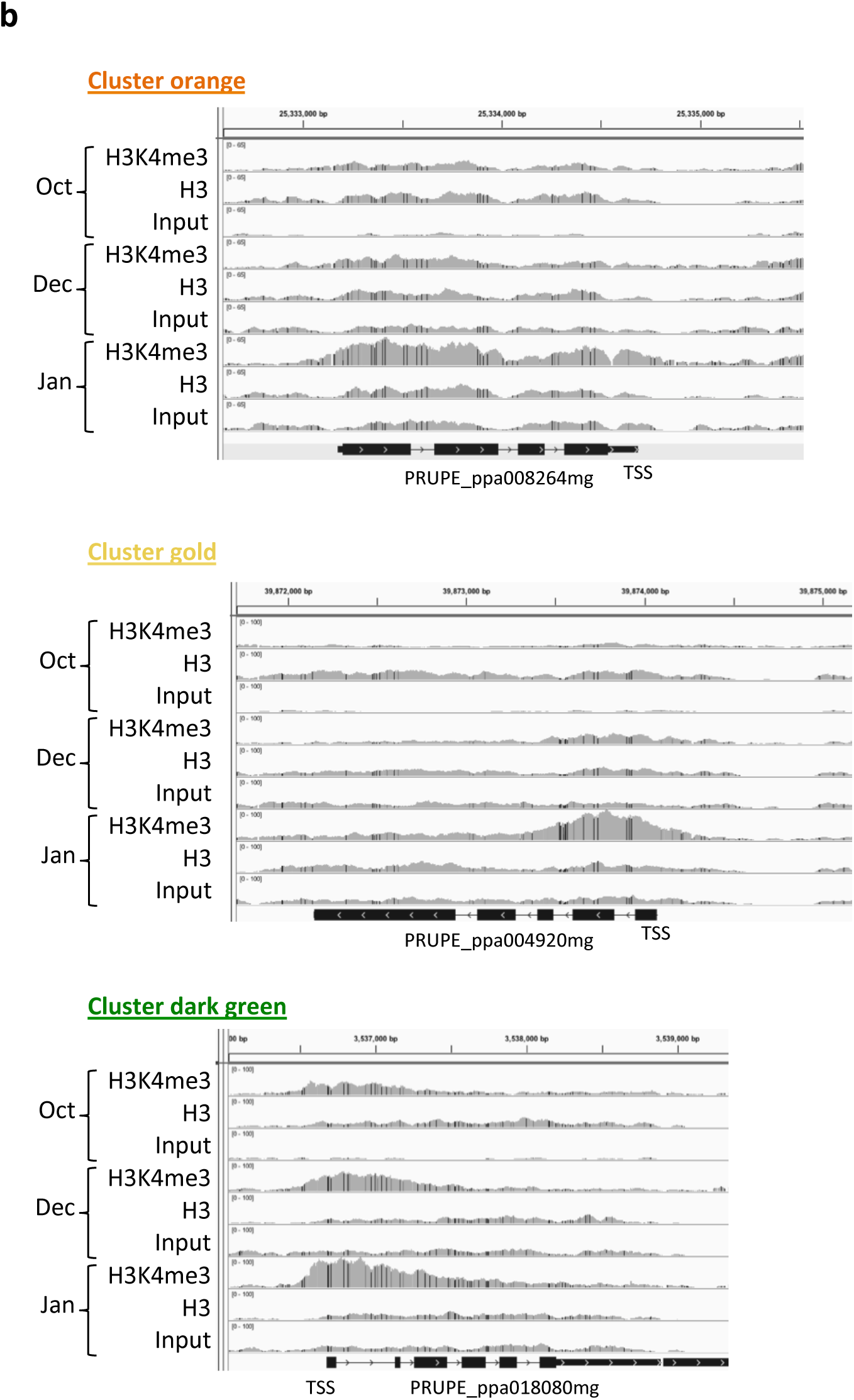
a) Average profile of H3K4me3 in the gene body of all genes, or around the TSS of all genes. Based on Cherry tree H3K4me3 ChIP-seq mapped in Peach genome. b) Example of H3K4me3 profile at the three dates for one gene in each of the six clusters.

